# Sexual deprivation induces a CRF independent stress response and decreases resistance to stressors in *Drosophila* via a subpopulation of Neuropeptide F receptor-expressing neurons

**DOI:** 10.1101/2022.03.02.482596

**Authors:** Julia Ryvkin, Anat Shmueli, Mali Levi, Avi Jacob, Tali Shalit, Assa Bentzur, Bella Agranovich, Ifat Abramovich, Eyal Gottlieb, Dick R. Nässel, Galit Shohat-Ophir

## Abstract

Living in a changing environment composed of other behaving animals entails both opportunities and challenges to obtain resources and mating partners. Actions that promote survival and reproduction are reinforced by the brain reward systems, whereas coping with the challenges associated with obtaining these rewards are mediated by stress response pathways. The activation of the latter can impair health and shorten lifespan. Although similar responses to social opportunity and challenge exist across the animal kingdom, little is known about the mechanisms that process reward and stress under different social conditions. Here, we studied the interplay between deprivation of sexual reward and stress response in *Drosophila melanogaster and* discovered that repeated failures to obtain sexual reward induces a frustration-like state that is characterized by increased arousal, persistent sexual motivation, and impaired ability to cope with starvation and oxidative stressors. We show that this increased arousal and sensitivity to starvation is mediated by disinhibition of neurons that express receptors for the fly homologue of neuropeptide Y (neuropeptide F, NPF). We furthermore demonstrate the existence of an anatomical overlap between stress and reward systems in the fly brain in the form of neurons that co-express receptors for NPF (NPFR) and the corticotropin-releasing factor (CRF)-like homologue Diuretic hormone 44 (Dh44), and that deprivation of sexual reward leads to translocation of forkhead box-subgroup O (FoxO) to the cytoplasm in these neurons. Nevertheless, the activity of Dh44 neurons alone does not mediate sensitivity to starvation and aroused behavior following sexual deprivation, instead, these responses are mediated by disinhibition of ~12-16 NPFR-expressing neurons via a dynamin-independent synaptic signaling mechanism, suggesting the existence of a NPFR mediated stress pathway which is Dh44-independent. This paves the path for using simple model organisms to dissect mechanisms behind anticipation of reward, and more specifically, to determine what happens when expectations to obtain natural and drug rewards are not met.

## Introduction

Living in a social environment involves diverse interactions between members of the same species, the outcomes of which affects health, survival, and reproductive success^1,2^. Coping with the challenges and opportunities associated with this dynamic environment requires individuals to rapidly process multiple sensory inputs, integrate this information with their own internal state and respond appropriately to various social encounters^2–8^. Encounters that hold opportunities to obtain resources, mating partners, and higher social status are considered rewarding and are therefore reinforced by the brain reward systems^9^, whereas the failure to obtain such rewards due to high competition or lack of competence can be perceived as a frustrative experience^10–16^. Deprivation or omission of expected reward (OER) occurs when organisms fail to obtain a reward they expect to receive despite signals for its presentation^10^. This condition leads to a frustrative state, characterized by increased motivation to obtain the reward^11,13,17^, increased levels of arousal^17^, agitation^12^, grooming^10,18^, stress associated behaviors^11,12^, drug consumption^13,19^, locomotion^12^, and aggression^13–16,20^. Example for a frustration-like state associated with deprivation of reward can be seen in small, submissive individuals of the rainbow trout, *Oncorhynchus mykiss,* which display an increase in both aggressive behavior and serotonin levels following frustrative reward omission, resulting in some cases in improved social status^20^.

Although certain stress responses can increase motivation and improve chances for obtaining natural rewards^21^, some types of social challenges like repeated aggressive encounters, lead to compromised health and shorten lifespan^22–28^. There is considerable anatomical overlap between brain regions that process reward and stress stimuli, as well as opposing functionality: exposure to natural rewards buffers the effect of stressors, while stressors such as social defeat can alter sensitivity to reward and increase the rewarding value of certain addictive drugs^2,22,22–27^. Example for such opposing functions is seen in rodents where Neuropeptide Y (NPY) decreases, while corticotropin-releasing factor (CRF) increases alcohol intake ^35–37^, and binding of NPY to NPY receptor Y1 that is found on CRF positive neurons within the bed nucleus of the stria terminalis (BNST) function to inhibit binge alcohol drinking ^29,37,38^.

Similar responses to social stress and reward-seeking behaviors can be seen in a variety of animals, suggesting that the central systems facilitating survival and reproduction originated early in evolution, and that similar ancient basic building blocks, biological processes and genes are inherently involved in these processes^7,39^. In agreement with this concept, we and others showed that *Drosophila melanogaster* display the ability to adjust their behavior and physiology to various changes in their social environment^40–52^. Moreover, recent studies provided evidence that the brains of mammals and fruit flies share similar principles when it comes to encoding stress and reward^40,53–55^. The homologs for NPY and CRF in Drosophila are Neuropeptide F (NPF) and Diuretic hormone 44 (Dh44), respectively^56–58^. Their effects on behavior are similar to those of mammals: the NPF and the NPF receptor (NPFR) system in flies mediates the response to reward and rewardseeking behavior, feeding behavior, suppresses responses to aversive stimuli such as harsh physical environments, decreases aggressive behaviors, and mediates courtship behavior^40,56,59–68^. Like CRF neurons, the Dh44 signaling pathway facilitates aggressive behavior in males, although whether Dh44 neurons mediate social stress response is still unknown^69^. Additionally, NPY\Y1 signaling is an essential link in a variety of processes affecting health and lifespan in mammals^70–74^, and similarly, activation of NPF neurons in male flies leads to decreased resistance to starvation and decreased lifespan^75,76^.

We previously showed that successful mating, and more specifically, ejaculation, is rewarding for male fruit flies^77^. Successful mating and sexual deprivation bi-directionally regulate the level of NPF, which in turn regulates the motivation to consume ethanol as a drug reward^40,78^. We recently discovered that male flies that experience repeated events of sexual deprivation respond to this social challenge by increasing their competitive behaviors in the form of increased arousal when interacting with rival male flies in a group, elevated aggression, and prolonged copulation time with receptive female flies (known as mate guarding)^47^. The latter is augmented by increased expression of certain seminal fluid genes that facilitate stronger postmating responses in female flies^47^. While certain types of social challenges in mammals are considered stressors, affecting health and lifespan, it is not known whether this occurs in fruit flies, and more specifically, whether failure to mate in *Drosophila* male flies is an acute stressor imposing certain costs to male flies, or simply a lack of reward.

In this study we show that repeated events of sexual deprivation in males lead to a frustration-like stress response, which is characterized by increased arousal, high motivational state and lower ability to endure metabolic and oxidative stressors. We then demonstrate the existence of an anatomical link between stress and reward systems within the fly brain in the form of receptors for NPF on Dh44 expressing neurons. Finally, we provide evidence that increased arousal and sensitivity to starvation stress result from disinhibition of a small subset of NPF-receptor neurons, which do not express Dh44 or NPF, and that this is mediated by a dynamin-independent, possibly peptidergic signaling mechanism.

## Results

### Sexual deprivation induces an acute stress response in male flies

The negative valence associated with courting non-receptive females and the behaviors exhibited by sexually deprived male flies prompted us to use courtship suppression as a tool to investigate the connection between deprivation of sexual reward and stress response in *Drosophila.* To determine whether deprivation/omission of sexual reward is experienced as stress, we closely observed the behavior of male flies during repeated encounters with previously mated non-receptive female flies. Wild type male flies were exposed to three consecutive one-hour interactions with mated females, spaced by one-hour rest (the ‘rejected’ cohort). The control cohort (naïve) consisted of naïve males, which were replaced at the beginning of each new session, and each time were coupled with new virgin females (Fig. 1A). The behavior of both cohorts during the first 10 min of the interaction was manually analyzed. In line with previous studies, the rejected cohort displayed a marked courtship suppression, reflected by a reduction in the overall time spent courting in all 3 sessions (Fig. 1B)^45,79,80^. Among those that exhibited active courtship during the sessions, various aspects of their courtship actions, such as wing vibration, licking and attempts to copulate were analyzed (Fig. 1B). To measure males’ innate response to first encounter with mated females, we analyzed all males in the first session. To discover which actions are affected by repeated encounter with mated females we analyzed the males that maintained active courtship during sessions 2 and 3.

**Fig 1.**
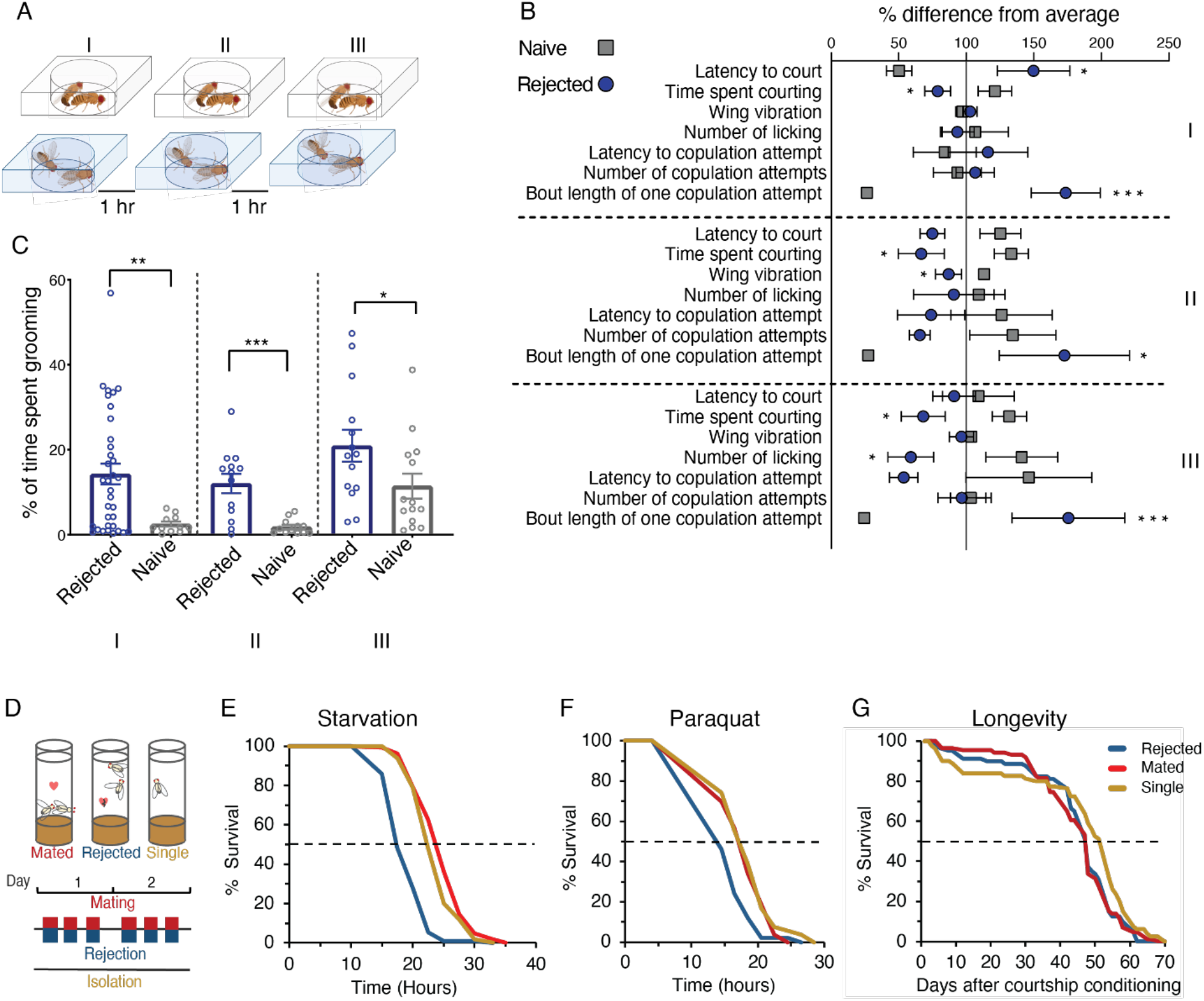
Repeated sexual deprivation increases sensitivity to stressors. **A.** Schematic representation of behavioral assays: Virgin male flies were exposed to either mated or virgin females for three 1h sessions, and their behavior was recorded. At the end of each session, the females and males from the naive cohort (top) were removed, and the males that experienced rejection (bottom) were kept isolated in narrow glass vials for 1h. **B.** % difference from average courtship behaviors performed by rejected (blue circle) and naive (grey square) males in the first (I), second (II), and third (III) sessions. Student’s t-test or Mann-Whitney were performed with FDR correction for multiple comparisons. \**p*<0.05, \*\*\**p*<0.001. **C.** % time spent grooming by rejected males (blue) compared to naive males (grey) during the first (I), second (II), and third (III) sessions. T-test or Mann-Whitney were performed. \**p*<0.05, \*\**p*<0.01, \*\*\**p*<0.001. N of rejected males= 41 (I),16 (II,III) (In order to capture differences in courtship behavior that resulted from a second and third exposure to mated females, only males that courted in both II, III sessions were analyzed). N of naive males= 24 (I), 21 (II), 18 (III). **D.** Schematic representation of courtship conditioning: naive males were introduced to either virgin, sexually receptive or sexually non-receptive females. As a result, males were either mated or rejected. The third cohort consisted of naive single housed males that did not experience any social or sexual event (single). Encounters with females were repeated 3x a day for two days. **E**. Starvation resistance assay: rejected males (blue, n=91) compared to mated (red, n=102) and single housed (yellow, n=110) males, *\*\*\*p*< 2E-16; mated vs single males, \**p*<0.05. **F.** Resistance to oxidative stress (20mM Paraquat): rejected males (blue, n=50) compared to mated (red, n=53) *p*<0.01** and single housed (yellow, n=54) males \*\*\**p*<0.001. Pairwise log-rank with FDR correction for multiple comparisons was performed in E,F. **G.** Longevity assay: single males (yellow, n=80) compared to mated (red, n=86) and rejected (blue, n=80) males. \**p*<0.05. Renyi-type test with PDF corrections for multiple comparisons was performed.

While the total percentage of time males spent courting was lower among rejected males compared to naive males in all three sessions (Fig. 1B), certain aspects of their courtship behavior were no less vigorous, such as the overall number of licking actions and attempts to copulate (Fig. 1B). Notably, although rejected males depicted longer latency to court mated females during the first encounter (consistent with innate aversion to the male pheromone cVA^79,81^), they overcame this aversion in subsequent sessions and initiated courtship at the same time as males that courted virgin females (Fig. 1B). Another feature that may reflect their surprising persistence was documented in the duration of copulation attempts, which was 6 times longer compared to controls (Fig. 1B), suggestive of persistent motivation to obtain mating reward. To our surprise, while rejected males depicted reduced courtship during the first 10 minutes of the interaction with mated females in courtship arenas (Fig. 1B), in a complimentary experiment, they did exhibit persistent courtship towards females along the entire training session, with no reduction in the overall number of males that exhibited courtship, during two entire days of consecutive rejection sessions (Fig. S1). These results suggest that rejected males experience a conflict between high motivation to mate and repeated inability to fulfil this mating drive.

We previously documented increased aggression, increased social avoidance and longer mating duration (LMD) following repeated sexual deprivation^47^. These actions can be interpreted as a result of a frustration-like state caused by stressful encounters with unreceptive females. To support this hypothesis, we searched for stress/frustration related actions that occur whenever animals are prevented from properly expressing a goal driven behavior. In such cases, rather than suspending actions, animals perform actions that are considered irrelevant to the context in which they take place (i.e. displacement behaviors) such as grooming, which is typically observed in stressful social situations^82–85^. In agreement with this, we documented a significantly higher incidence of self-grooming in rejected males in all 3 sessions, suggesting that selfgrooming is a displacement-like action that is associated with the failure to mate (Fig. 1C).

In mice, acute social stress caused by aggressive encounters with a dominant male is known to shorten lifespan and reduce the ability to cope with other types of stressors^24–28^. The appearance of displacement like behavior, increased aggression, and elevated arousal prompted us to further test the hypothesis that deprivation of sexual reward is perceived as a stressor to male flies, which can impair their ability to cope with additional stressors. To test this, three cohorts of male flies were exposed to the following social conditions: (a) males that experienced multiple events of successful mating encounters over the course of two days (mated cohort), (b) males that experienced multiple events of sexual deprivation by previously mated, non-receptive females (rejected cohort) and (c) males that were housed in isolation for the entire duration of the experiment (single cohort) (Fig. 1D). Each cohort was then exposed to starvation (1% agarose) or oxidative stress (20mM paraquat), and the rate of survival over time was documented (Fig. 1D-F). If deprivation of sexual reward is a stressor, we expect the cohort of rejected male flies to be more sensitive to other stressors. Indeed, rejected males exhibited higher sensitivity to both starvation and oxidative stress compared to controls. While mated and single cohorts reached 50% decline in survival after 22-24h of starvation, rejected male flies exhibited a faster decline in survival rates, reaching 50% survival after less than 18h (Fig. 1E). Exposure to paraquat led to a 50% decline in survival after more than 17h for both the mated and single cohorts, vs. less than 14 hours for rejected males (Fig. 1F). This indicates that deprivation of sexual reward promotes sensitivity to starvation and oxidative stress, further suggesting that multiple rejection events are perceived as an acute stressor, compromising the ability of male flies to cope with other stressors. Extending this line of experiments, we next examined whether deprivation of sexual reward also affects the overall lifespan of rejected male flies, by quantifying the longevity of the 3 cohorts of flies over the course of 70 days. While rejected and mated cohorts exhibited shorter lifespans compared to isolated male flies, there were no significant differences between rejected and mated cohorts (Fig. 1G). The prolonged lifespan of single male flies suggests that any interaction with female flies is sufficient to reduce the longevity of male flies (even as little as 6h), and is in agreement with previous studies showing that exposure to female pheromones shortens the lifespan of male flies^75,86^. The lack of differences between the rejected and mated cohorts suggests that deprivation of sexual reward does not affect general lifespan, but rather affect the way by which male flies cope with additional acute stressors.

### Deprivation of sexual reward is represented by discrete transcriptional programs in NPFR neurons

To explore mechanisms that could explain the apparent trade-off between the behavioral responses to being repeatedly rejected and the sensitivity to subsequent stressors, an unbiased transcriptomic approach was undertaken to highlight changes in brain transcriptional programs in response to deprivation of sexual reward. For that, we isolated and sequenced the transcriptome of a selected neuronal population from brains of male flies that experienced either two days of repeated rejection events, successful mating, or single housing. Celltype specific RNA isolation was achieved using the INTACT (Isolation of Nuclei Tagged in specific Cell Types) method, which utilizes immunoprecipitation of genetically marked nuclei (Fig. 2A)^87^. We chose to isolate NPF-receptor expressing neurons based on their role in processing sexual interactions, motivational drives, reward seeking behavior and aggressive and aroused behaviors^40,62^ (Fig. 2B). Analyzing the transcriptome of the 3 different conditions revealed 258 differentially expressed genes (DEGs) with discrete transcriptional programs for rejection, successful mating, and single housing (Fig. 2C, Table S1).

**Fig 2.**
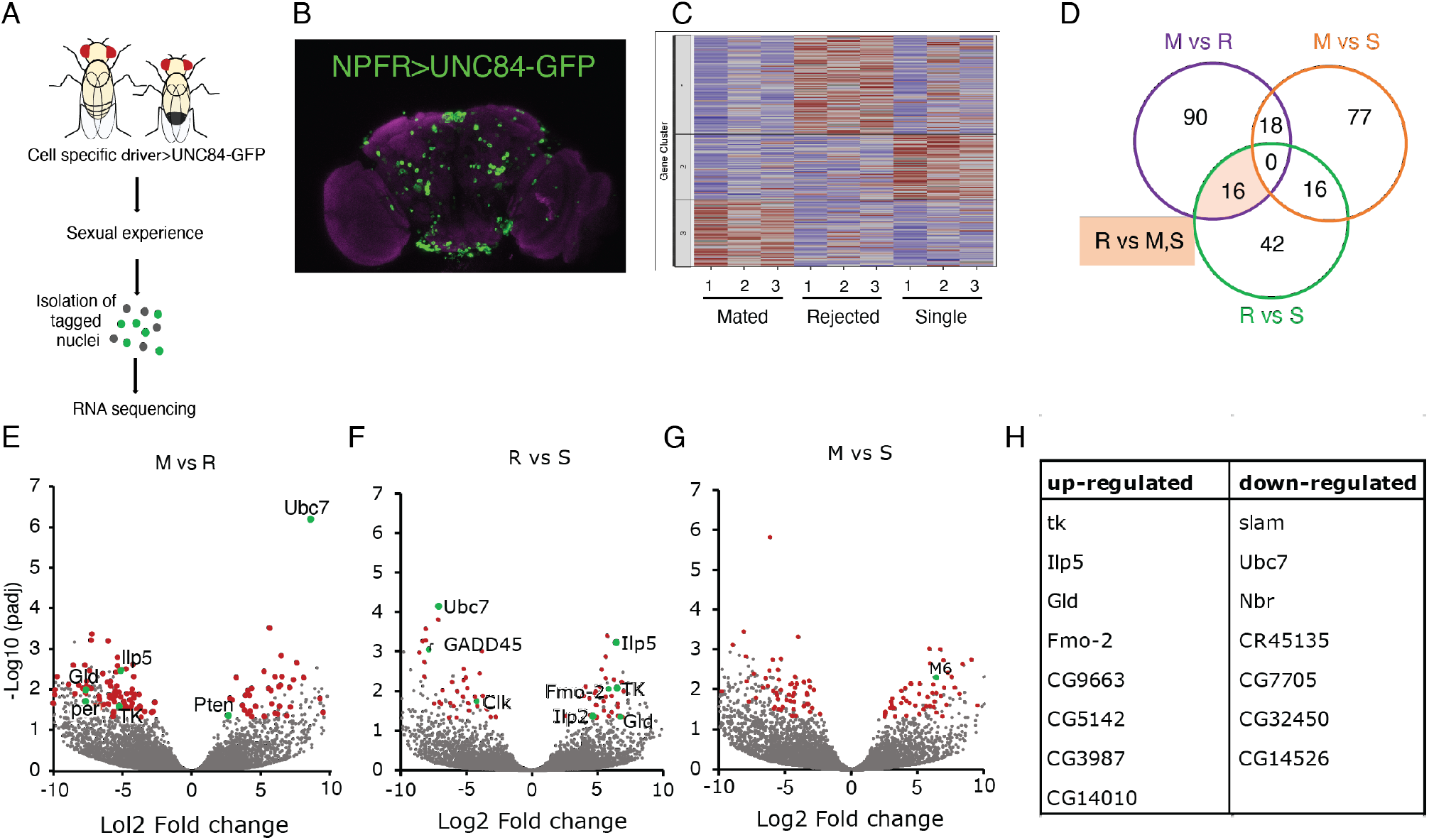
Courtship conditioning induces differential gene expression for rejected, mated, and single housed males. **A**. Schematic representation of INTACT procedure. **B.** Distribution of NPFR (Green) neurons as visualized by the expression of GFP driven by NPFR-GAL4 driver. Anti-nc-82 staining is shown in magenta. **C.** Clustering of average normalized reads for all significantly differentially expressed genes in NPFR neurons of rejected, mated, and single housed males. **D**. Venn diagram depicting differentially expressed genes shared among the courtship conditioning cohorts. Highlighted in beige are genes that were differentially expressed in rejected males compared to mated and single males. **E-G.** Volcano plots of genes that were differentially expressed when comparing: Mated vs rejected (**E**), rejected vs single (**F**) and mated vs single (**G**). **H.** Table of genes up-regulated (left) and down-regulated (right) in rejected males compared to mated and single males.

We documented 77 DEGs in mated vs. single male flies, 90 DEGs in mated vs. rejected male flies, and 42 DEGs when comparing rejected to single males (Figure 2D, Table S2). Analyzing the statistical overrepresentation of known biological processes of all DEGs using the PANTHER GO-term analysis, did not reveal overrepresentation of specific GO term groups indicative of common pathway or cellular function. Nevertheless, the identified DEGs can shed light on gene products that function in shaping behavioral and physiological responses to changes in social conditions. Interesting examples are two circadian clock genes, Clock (Clk) and period (per), in rejected males (Fig. 2E-F). Tachykinin (tk), a neuropeptide that functions to regulate aggressive arousal and process anti-aphrodisiac pheromone information was upregulated in response to rejection (Fig. 2E,F,H)^41,51,88^. Another courtship-related gene identified in our dataset was Ubiquitin-conjugating enzyme7 (Ubc7), which was down-regulated in response to rejection (Fig. 2E,F,H)^60,89^. Focusing on DEGs specific for the rejected cohort, we identified 15 genes (Fig. 2H) that were enriched or depleted in rejected males compared to both single and mated flies. Manual annotation of these indicated a few genes associated with the metabolic response to starvation and oxidative stress or related to the insulin/FoxO stress response pathway, such as Insulin-like peptide 5 (Ilp5), Flavin-containing monooxygenase 2 (Fmo2) and Glucose dehydrogenase (Gld) (Fig. 2H)^90–95^. This suggests a molecular connection between metabolic\cellular stress response pathways and the experience of deprivation of sexual reward.

### Sensitivity to starvation and oxidative stress is not a result of cytoprotective activity in DH44 positive NPFR neurons

Expression differences in genes associated with metabolism and stress response following rejection support two possible explanations for what appears to be a trade-off between short-term behavioral responses and higher sensitivity to starvation and oxidative stress. The first is that enhancement of some courtship behaviors during rejection generates a high metabolic demand, resulting in depletion of reservoirs and therefore an increase in sensitivity to starvation. The second is that the challenge of repeated rejection and the resulting frustration-like state induces oxidative stress or activates a neuronal stress response that impairs the cytoprotective response to starvation. To test whether rejected male flies are subjected to high energetic demand, we analyzed triglycerides (TAG), fly weight and body or hemolymph glucose levels in rejected, mated and single male flies. No difference was documented in body TAG, glucose levels, or body weight between the cohorts (Fig. 3A-D), suggesting that rejected male flies do not suffer from an energetic deficit which would increase their sensitivity to starvation.

**Figure 3.**
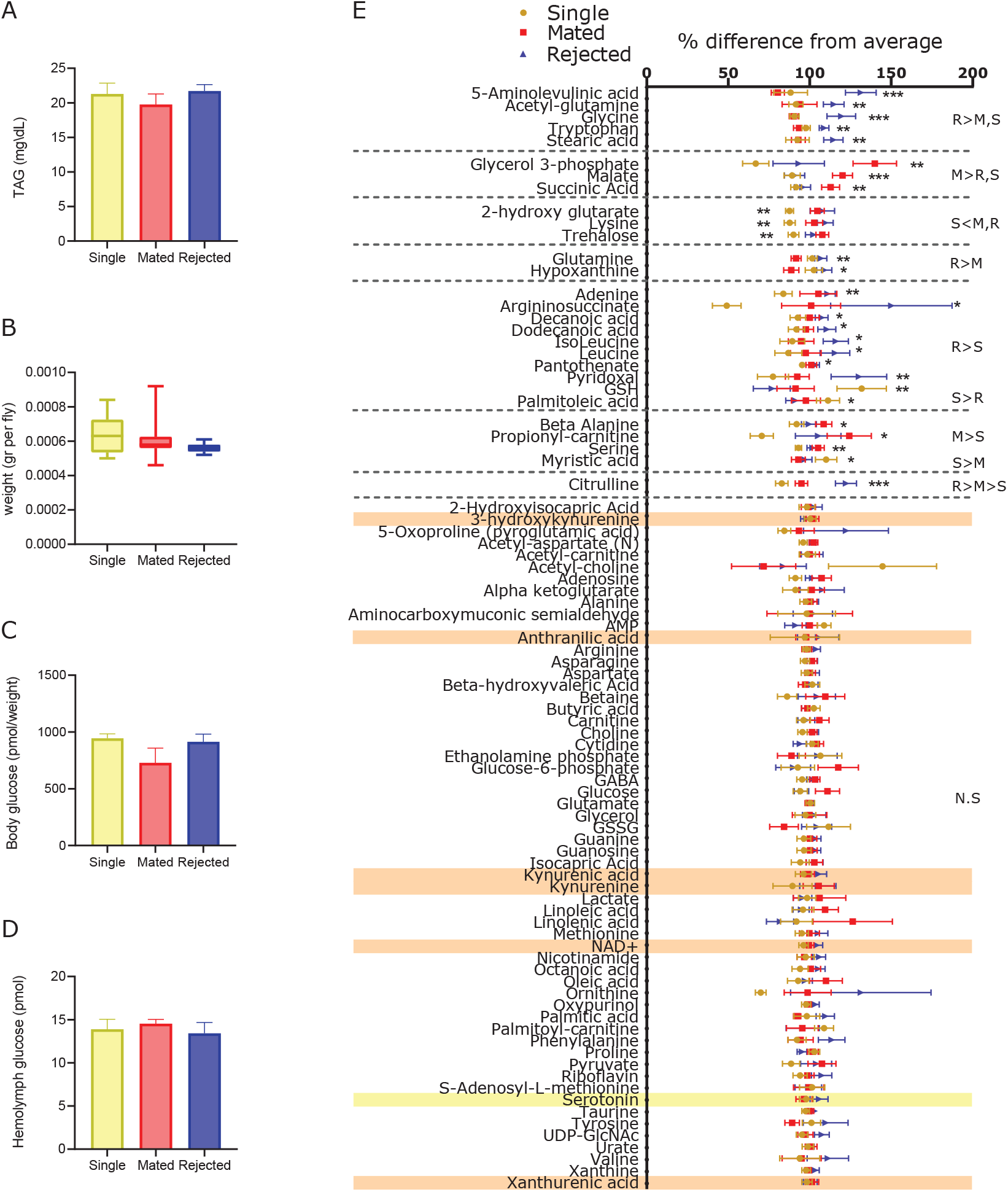
Courtship conditioning did not affect TAG and glucose levels and most head metabolite in males. **A-D.** Metabolic indices of rejected males (blue) compared to single (yellow) or mated (red) males. No differences were observed for measurements of (**A**) triglycerides (TAG, n=11 for all groups, 5 males/ sample). (**B**) weight (n= 10 single, 9 mated, 9 rejected, 5 males/sample). (**C**) hemolymph, or (**D**) body glucose (n=3 for all groups, 5 males/body sample, and 40 males/hemolymph sample). ANOVA or Kruskal-Wallis with post-hoc Tukey’s or Friedman test were performed. NS *p*>0.05. **E.** % difference from average (peak area/ total measurable ions) of metabolites detected using LC-MS in rejected, mated, and single males’ heads. 5-aminolevulinic acid, acetyl-glutamine, glycine, tryptophan, and stearic acid levels were higher in rejected males’ heads (blue triangles, n=17) compared to single (yellow circles, n=17) and mated (red squares, n=16, 5 heads/sample). Metabolites of the kynurenine pathways are highlighted in orange; serotonin is highlighted in yellow. NS *p*>0.05, \**p*<0.05, \*\**p*<0.01, \*\*\**p*<0.001. Statistical analysis was performed by ANOVA or Kruskal-Wallis with post-hoc Tukey’s or Friedman test.

The brain, the central organ that controls stress response, is structurally and chemically sensitive to stress, especially oxidative stress^96^. We hypothesized that certain brain metabolites are produced differentially in response to acute social stress, affecting starvation and oxidative stress resistance. To capture metabolic changes, which could potentially affect resistance to stressors, we extracted head metabolites of males that experienced courtship conditioning using polar solvents and performed liquid chromatography-mass spectrometry (LC-MS) for targeted metabolite analysis. Analysis of the metabolome revealed a unique profile for each cohort (Fig. 3E). Looking only at metabolites which were enriched or depleted in rejected compared to both mated and single cohorts, five metabolites were enriched in rejected males: Glycine, tryptophan, 5-Aminolevulinic acid (5ALA), acetyl-glutamine, and stearic acid (Fig. 3E, Table S3). Though tryptophan can be converted to serotonin or tryptamine, and most of the tryptophan metabolizes to kynurenine pathway (KP) metabolites, which contribute to a shorter lifespan^97–100^, we did not document significant accumulation or decline of metabolites in the KP (Fig. 3E). Additionally, we did not document a difference in the abundance of serotonin (Fig. 3E). No difference was observed in oxidative agents nor in antioxidant activity and glucose levels, though a decline in trehalose (the main circulating sugar in flies) levels was detected in single males compared to mated and rejected males, indicating a possible change in insulin signaling^101–104^. In summary, no significant changes in the metabolome were detected that could explain the enhanced sensitivity to stressors observed in rejected flies, suggesting this is not a result of metabolic effects.

Activation of the insulin signaling pathway can decrease lifespan and affect resistance to starvation, mainly via FoxO phosphorylation^105–111^. The connection between FoxO activity and lifespan\resistance to stressors has been studied in *Drosophila,* typically in the fat body, where the activity of FoxO can prolong lifespan and increase resistance to stressors^105,112^. FoxO activity in abdominal and\or pericerebral fat bodies affects insulin signaling, lipolysis, gluconeogenesis and thus the abundance of TAG and different metabolites^86,92,105,112–114^. The lack of changes in TAG, glucose, and oxidative stress related metabolites in response to deprivation of sexual reward indicates that FoxO activity in fat body is not affected, and instead suggests a potential role in neurons, as supported by our RNAseq results. To test this possibility, we examined expression pattern of endogenous FoxO in the brain using FoxO specific antibodies. In agreement with Cao et al^115^ and Okamoto, et al.^88^, we identified six FoxO positive neurons in the pars intercerebralis neurons (PI) (Fig. 4A). These FoxO expressing neurons were colocalized with NPFR expressing insulin-producing cells (IPCs) that could be identified by Ilp2 immunolabeling (Fig. 4A, S2A,B). Based on similarity in the transcriptional profile of Dh44 and NPFR neuronal populations identified in our previous study^60^, we examined the overlap between the two populations as well as FoxO expression. Indeed, all six PI Dh44 neurons are NPFR (Fig. 4B), and five of them also express FoxO (Fig. 4C). This indicates that these are NPFR-Dh44+ and FoxO expressing neurons.

**Fig 4.**
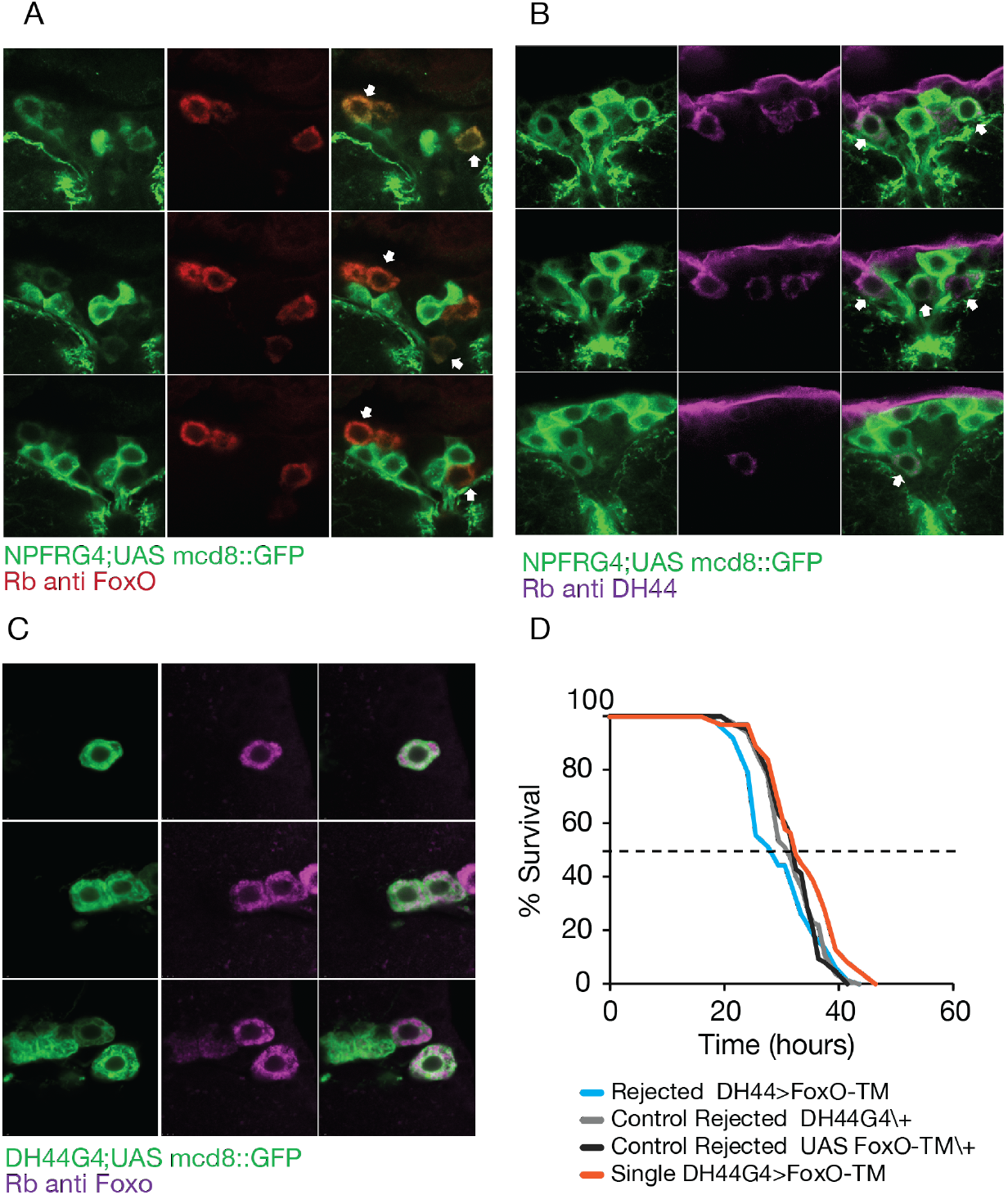
FoxO activity in NPFR-Dh44 neurons does not abrogate sensitivity to starvation in rejected males. **A.** Six NPFRG4; UAS-mcd8GFP (green) neurons co-localized with FoxO (red, endogenous FoxO expression), as indicated by arrows. **B.** Co-localization of NPFR neurons (green, NPFRG4/+;UAS-mcd8GFP/+) with six Dh44 neurons (magenta, endogenous Dh44 expression). **C.** At least five Dh44 neurons (green, Dh44 G4/+;UAS-mcd8GFP/+) co-localized with FoxO+ neurons (magenta, endogenous expression). **D.** Sensitivity to starvation was assessed in rejected males with UAS-FoxO-TM expressed in DH44 neurons (DH44G4;UAS-dFoxO-TM, blue), and their genetic controls (light and dark grey), compared to single housed males (red) (\**p*<0.05). NS, not statistically significant (*p*>0.05). Pairwise log-rank with FDR correction for multiple comparisons was performed.

Next, we examined whether sexual deprivation inactivates FoxO by forcing its translocation to the cytoplasm of neurons, thereby preventing it from inducing transcription of genes important for survival. We imaged the subcellular localization of FoxO in brains of male flies that experienced sexual deprivation, mating or were single housed at the end of the conditioning phase or after 20 hours of starvation. The mean fluorescence intensity of FoxO within the cytoplasm (cyto) was normalized to the mean fluorescence intensity of FoxO in the nucleus (cyto/nuc) and compared among rejected, mated, and single housed males. Flies which were not subjected to starvation showed no significant difference in mean fluorescence intensity between the cohorts (Fig. S3A-D), whereas starved cohorts exhibited significant differences between conditions. Rejected males showed a higher relative cyto/nuc intensity of fluorescence compared to both mated and single males (Fig. S3E-H), and both rejected and mated males displayed higher relative mean cyto/nuc intensity of fluorescence than single housed males, perhaps due to exposure to female pheromones (Fig. S3E-H). This suggests that sexual rejection changes FoxO translocation to the nucleus, which might affect sensitivity to stressors.

To test whether export of FoxO from the nucleus is required for the increased sensitivity of rejected males to starvation, a form of FoxO that is mutated in three phosphorylation sites such that it is retained in the nucleus (UAS-dFoxO-TM)^105^ was expressed in Dh44 neurons. Dh44> dFoxO-TM flies were exposed to repeated sexual deprivation and their sensitivity to starvation stress was analyzed. While, rejected males expressing nuclear FoxO exhibited similar response to starvation as their corresponding genetic controls, they were more sensitive to starvation than Dh44> dFoxO-TM single housed males flies, similar to the differences between rejected and single housed observed in WT males (Fig. 1E, Fig. 4D). Altogether, these results suggest that repeated sexual deprivation, followed by starvation affects FoxO localization in Dh44 neurons. However, FoxO activity in these neurons does not directly facilitate sensitivity to starvation in response to sexual deprivation.

### Disinhibition of NPFR neurons increases sensitivity to starvation in male flies

Given the bidirectional regulation of NPF levels in response to sexual deprivation and mating, where deprivation reduces and mating induces its levels^40^, and the inhibitory effect of NPY binding to NPY receptor neurons^29,56,116–118^, we postulated that the disinhibition of NPFR neurons in response to sexual deprivation promotes sensitivity to starvation. To test this hypothesis, we knocked down NPFR in NPFR neurons and compared the sensitivity of single, rejected and mated male flies to starvation stress. If sexual deprivation reduces NPF signaling, that in turn disinhibits NPFR neurons, we expect mated male flies to exhibit similar responses as rejected male flies. This manipulation abrogated the differences between mated and rejected cohorts (Fig. 5A), implying that NPFR is necessary for this effect. Interestingly, NPFR KD in single housed males showed higher resistance to starvation compared to both rejected and mated males (Fig. 5A), implying that NPFR activity has no effect on single housed males. It also emphasizes that NPFR KD increased mated males’ sensitivity to starvation rather than reducing rejected males’ sensitivity. This suggests that the activity of NPFR neurons mediates the perception of non-rewarding experience in the context of a sexual experience, and consequently increases sensitivity to starvation.

**Fig 5.**
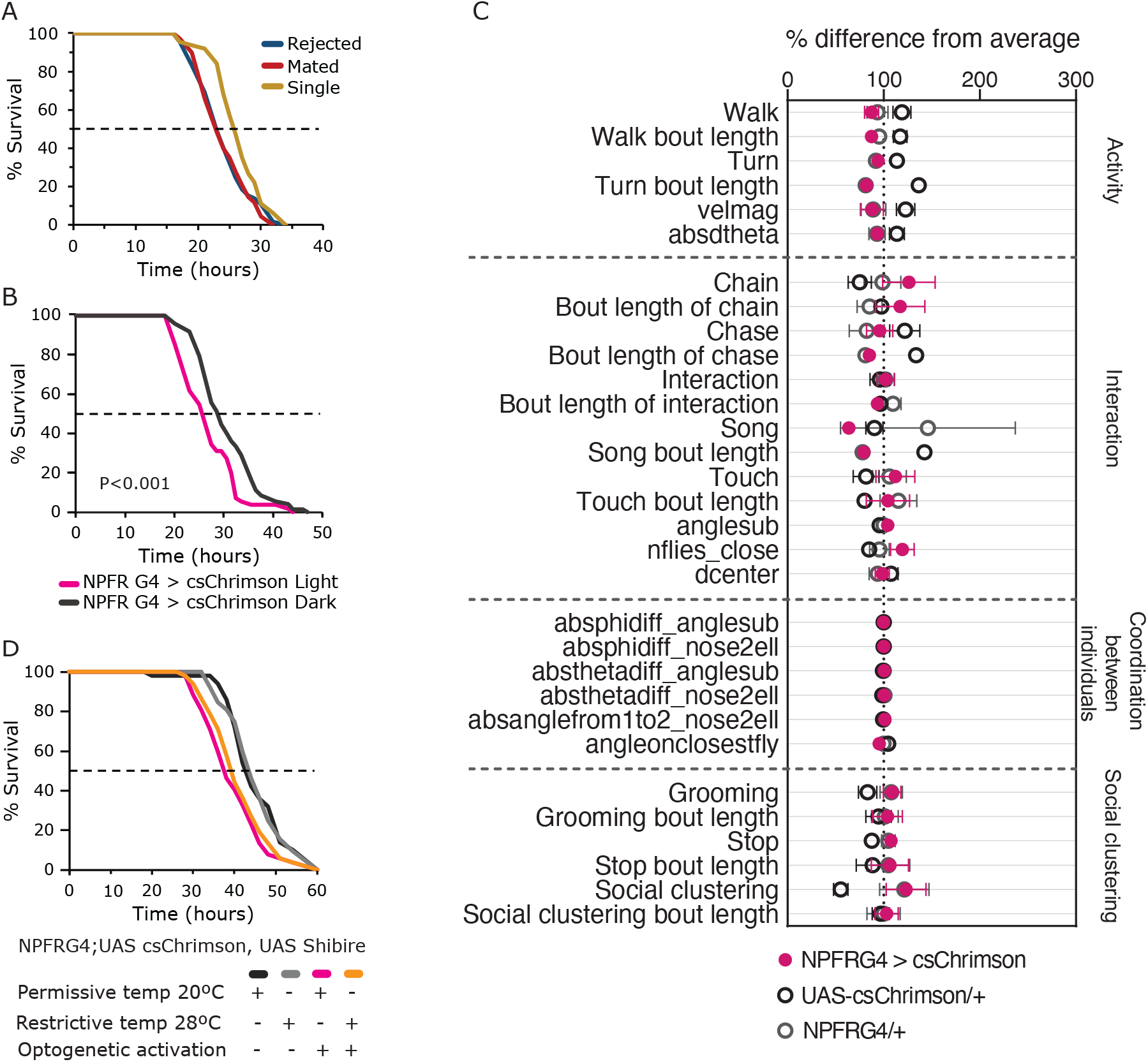
Activation of NPFR neurons mimics reduced resistance to starvation observed in WT males. **A.** Starvation resistance assayed on NPFR G4/NPFR RNAi mated (red, n=68), rejected (blue, n=68) and single housed (yellow, n=63) male flies. No significant difference was observed among mated and rejected flies *p*>0.05. Both rejected and mated males were significantly more sensitive to starvation compared to single males. \**p*<0.05 rejected vs single, \*\**p*<0.01 mated vs single. Pairwise log-rank with FDR correction for multiple comparisons was performed. **B.** NPFR neurons were activated in NPFRG4/+;UAS-csChrimson/+ flies (pink) by exposing them to red light three times a day for two days, while control flies (grey) were not exposed to light. Starvation resistance of experimental (n=55) and control flies (n=72) was assayed. Log-rank test was performed, \*\*\**p*< 0.001. **C.** % difference from average scatter plot of behaviors observed and scored in the FlyBowl performed by NPFRG4/+;UAS-csChrimson/+ males (pink, n=14) and their genetic controls NPFR G4/+, UAS-csChrimson/+ (Grey and black, respectively, n=10 each). ANOVA or Kruskal-Wallis with post-hoc Tukey’s or Dunn’s test with FDR correction for multiple comparisons was performed. **D.** Male flies expressing UAS-csChrimson and UAS-Shibire^ts^ in NPFR neurons were subjected to three 5 min long optogenetic activations for three days, and their synaptic signaling was blocked at 28-29°C (light+ heat, orange, n=52). Positive control males (light+cold, pink, n=52), synaptic release block control (dark+heat, light gray, n=52), negative control (dark+cold, dark gray, n=50). Experimental and positive control flies showed no significant difference in resistance to starvation (*p*>0.05). Both experimental and positive control flies were significantly more sensitive to starvation than ‘dark+heat’, and ‘dark+cold’ flies (\*\**p*<0.01). Pairwise log-rank test with FDR correction for multiple comparisons was performed.

To further strengthen this finding and mimic rejection state, we optogenetically activated NPFR neurons in naïve NPFR>csChrimson male flies by repeatedly exposing them to red light for two days, three times each day, and tested their resistance to starvation. Experimental flies were significantly more sensitive to starvation than controls (Fig. 5B), implying that activation of NPFR neurons is sufficient to mimic the effect of sexual deprivation on sensitivity to starvation. To assess whether activation of these neurons also mediates other behavioral phenotypes observed in rejected males, we assayed the behavior of NPFR>csChrimson flies during optogenetic activation in an open field exploration arena (FlyBowl) in which flies can move and interact in two dimensions. The system is coupled with a tracking and machine learning algorithms capable of automatically quantifying various behavioral parameters^60,119^. Intriguingly, we observed no difference in the behavior of experimental flies compared to both genetic controls (NPFRG4\+, UAS-csChrimson\+) (Fig. 5C, S4), suggesting that disinhibition of all NPFR neurons is sufficient to cause increased sensitivity to starvation, but not to induce changes in social interaction.

We next examined whether synaptic signaling is required for starvation sensitivity that is triggered by activation of NPFR neurons. To test this, NPFR neurons were optogenetically activated while their synaptic transmission was blocked using temperature-sensitive Shibire (Shibire^ts^). Activation of NPFR neurons and simultaneous inhibition of their synaptic transmission resulted in similar levels of sensitivity to starvation as that of activation of NPFR neurons alone (Fig. 5D), suggesting that sensitivity to starvation induced by sexual deprivation is not mediated via a synaptic dynamin-based neurotransmitter release, but possibly via peptidergic signaling. Since Insulin signaling is dynamin independent, we tested whether sexual deprivation or activation of NPFR neurons induces insulin release from IPCs. Imaging of endogenous Ilp2 in brains of males that experienced sexual deprivation or optogenetic activation of NPFR neurons identified no apparent difference in its abundance in under all tested conditions (Fig. S5A-E).

### Activation of a small subpopulation of NPFR neurons increases sensitivity to starvation and promotes social arousal

So far, we demonstrated that optogenetic activation of NPFR neurons is sufficient to recapitulate the increased sensitivity to starvation but does not mimic the increased arousal and frustration-like behavior associated with deprivation of sexual reward. Since the NPFR neuronal population consists of about 100 neurons, we next investigated which subset of NPFR cells is responsible for the enhanced sensitivity to starvation. Given the possibility that sensitivity to starvation is mediated via neuropeptide release, we divided the NPFR expression pattern into sub-populations using genetic intersection with drivers for neuropeptides that are co-expressed with NPFR such as NPF^120^, Dh44 and Tachykinin (illuminated in our RNAseq dataset, Fig. 2E,F,H, Table S2). Activating Dh44 neurons did not affect sensitivity to starvation and did not lead to apparent behavioral responses using the FlyBowl system (Fig. 6A, S6B). In addition, knocking down the expression of Dh44 in NPFR neurons did not change sensitivity to starvation (Fig. S6A), altogether suggesting that Dh44 signaling pathway does not mediate the sexual reward deprivation stress response. Similarly, activation of NPFR^NPF^ mutual cells (P1 and L1-1 neurons) did not affect either sensitivity to starvation or group behavior in the FlyBowl (Fig. 6B, S6C), suggesting that subpopulations of NPF expressing NPFR neurons, which presumably have the capacity for autoinhibition are not responsible for these effects.

**Fig 6.**
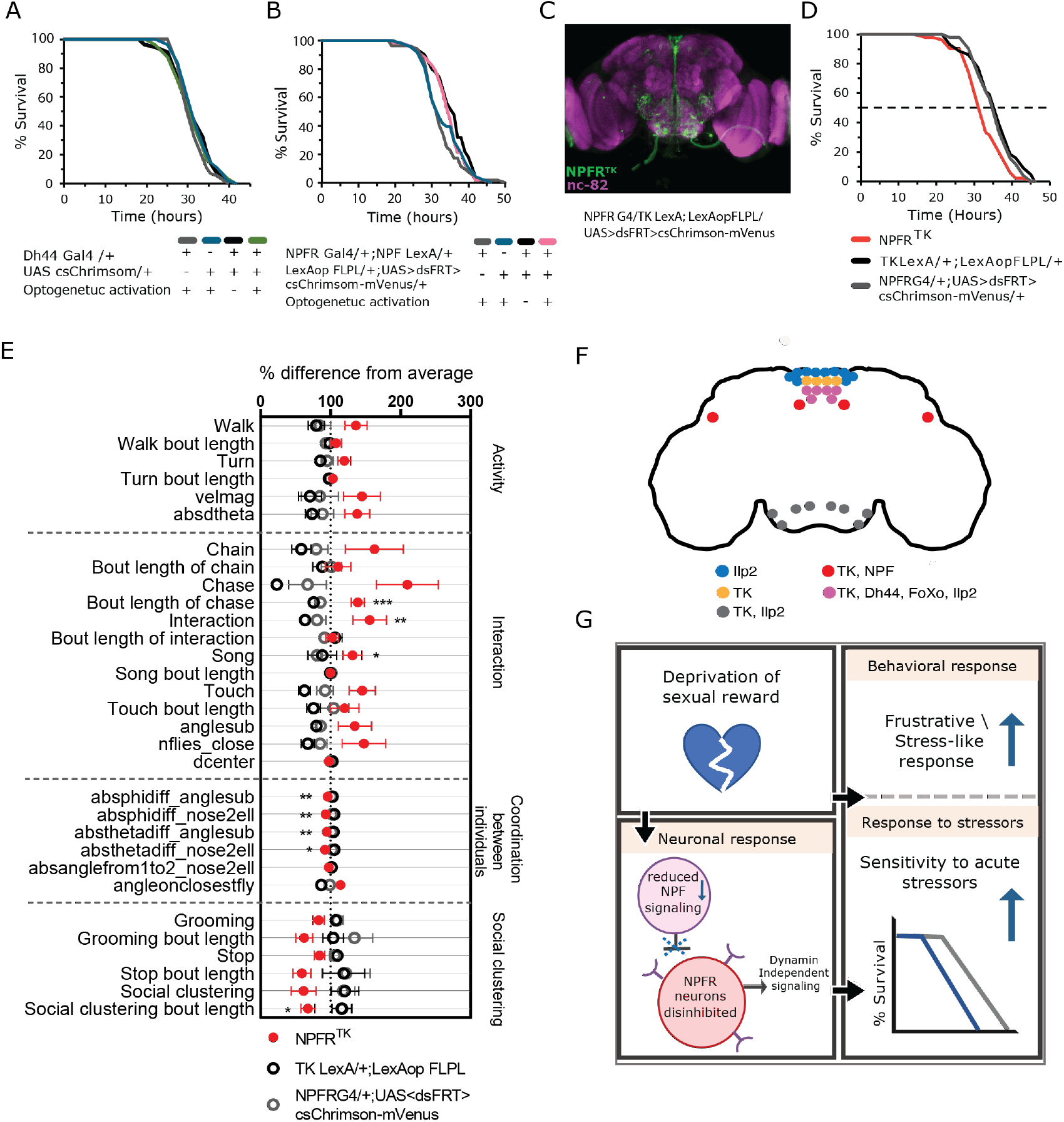
Activation of NPFR^TK^ neurons reduces resistance to starvation and causes aroused behavior. **A.** Dh44 neurons were activated in Dh44 G4/+;UAS-csChrimson/+ flies followed by starvation resistance assay. No significant difference was observed between experimental (Light, green), genetic controls (Dh44-G4/+, grey, UAS-csChrimsom/+, dark blue) and Dh44 G4/+;UAS-csChrimson/+ flies that were not exposed to light (Dark, black). *p*>0.05, n=56 for all cohorts. **B.** Intersection of NPFR with NPF neurons by crossing NPFRG4;+, NPF-LexA;+, +;LexAop-FlpL, +;UAS<dsFRT>cs-Chrimson-mVenus flies. Naïve experimental males (Light, pink), n=52 and their genetic controls (NPFR G4/+;NPF LexA/+, grey, n=52; LexAop-FLPL/+;UAS<dsFRT>cs-Chrimson-mVenus/+, dark blue, n=58) were exposed to red light three times a day for two days. NPFR-NPF flies that were not exposed to light served as a third control (Dark, black, n=50). Experimental flies (pink) did not exhibit significantly different resistance to starvation compared to control flies (grey) *p*>0.05. **C.** Intersection of NPFR neurons with TK neurons (NPFR^TK^) by crossing NPFRG4;+, TK-LexA;+, +;LexAop-FlpL, +;UAS<dsFRT>cs-Chrimson-mVenus flies. Green marks NPFR^TK^ neurons, magenta marks nc-82 neurons. **D.** NPFR^TK^ neurons of naive males were activated, and their starvation resistance was assayed. Experimental flies (NPFR^TK^, red, n=51) exhibited significantly lower resistance to starvation compared to genetic control flies (TK-LexA;LexAop-FLPL, black, n=44; and NPFR G4;UAS<dsFRT>csChrimson-mVenus, grey, n=51). \**p*<0.05, \*\**p*<0.01. Pairwise log-rank test with FDR correction for multiple comparisons was performed for A,B,D. **E.** Scatter plot of behaviors observed and scored in the FlyBowl performed by NPFR^TK^ males (red, n=13) and their genetic controls, TK-LexA; LexAop-FLPL, NPFRG4;UAS<dsFRT>csChrimson-mVenus (black, n=13, and grey, n=12. \**p*<0.05,\*\**p*<0.01, \*\*\**p*<0.001. ANOVA or Kruskal-Wallis with post-hoc Tukey’s or Dunn’s test, and FDR correction for multiple comparisons were performed. **F**. Schematic representation of brain NPFR neurons that intersect with TK, Ilp2, DH44, FoxO and NPF neurons. **G.** Summary of results: Deprivation of sexual reward induces a frustrative-like or a stress-like behavioral response and increases sensitivity to subsequent acute stressors. This is mediated by a neuronal response: Sexual deprivation decreases NPF signaling, thereby disinhibits NPFR neurons and induces a dynamin-independent activity, which increases sensitivity to starvation.

Finally, the activation of a subpopulation of 22-26 NPFR neurons that co-express the neuropeptide Tachykinin (NPFR^TK^) (Fig. 6C) induced starvation sensitivity that was similar in its extent to activation of the entire NPFR population (Fig. 6D). Knock down of Tachykinin in NPFR did not affect sensitivity to starvation (Fig. S7A,B), suggesting that the activity of NPFR^TK^ neurons and not Tachykinin neuropeptide mediate activation induced sensitivity to starvation. Next, we tested the behavioral responses associated with the activation of NPFR^TK^ neurons in the FlyBowl system. NPFR-TK>csChrimson males exhibited increased social arousal, characterized by higher levels of active approach behavior, increased duration of chase behavior, an increase in unilateral wing vibration (song), and in many cases, formed long chains containing 4-8 flies (Fig. 6E). Moreover, experimental flies depicted a significant reduction in features that reflect changes in angle and speed between two close individuals (absanglefrom 1to2, absphidiff, absthetadiff, and angleonclosestfly; see Fig. S4 for more details). This may signify an increase in coordination between pairs of flies and suggest that NPFR-TK>cs-Chrimson flies engage more persistently with others when interacting (Fig. 6E). Lastly, experimental flies exhibited reduced social clustering (Fig. 6E), altogether suggesting that activation of NPFR^TK^ neurons mimics some of the activity-related behavioral effects induced by sexual deprivation: reduced social clustering and overall arousal. Taken together, this implies that although expression of tk transcript in NPFR neurons does not affect sensitivity to starvation, disinhibition of NPFR^TK^ neurons does affect sensitivity to starvation and produces a behavioral phenotype which resembles a frustration\stress-like behavior. Analyzing the expression pattern of NPFR^TK^ neurons, we noticed that part of this neuronal population coexpresses Dh44 and NPF (Fig. 6F, S7C,D), suggesting that only about 12-16 NPFR^TK^ neurons that do not express Dh44 and NPF regulates both sensitivity to starvation and tunes arousal levels. This further indicates that stress-like response to reward deprivation is mediated by NPF/NPFR rather than a Dh44 circuit.

## Discussion

Reward-seeking behaviors are evolutionary adaptations designed to increase motivation to perform actions that in turn will increase fitness. When the expectation of reward is not met, individuals exhibit frustration and change their behavior towards other animals^10–16^. Here we utilized courtship suppression to deprive male flies of the inherent expectation of sexual reward as a model for frustration-like stress response in *Drosophila*. While courtship suppression is associated with reduced courtship and presumably a defeat-like state, we discovered that rejected males are rather persistent in their attempts to obtain sexual reward. This finding is not completely surprising considering the innate nature of mating motivation and the presence of female aphrodisiac pheromones. It can also be attributed to measuring various courtship actions rather than overall percentage of time spent courting, which is the usual indicator used to quantify courtship.

In response to repeated failures to mate, rejected males exhibit features characteristic of a frustration-like state, such as persistent mating actions (some of which are elongated), increased arousal, increased aggression, as well as long mating duration upon successful mating encounters, all of which are reminiscent of high motivational state^47^. While this motivational state may serve to increase their fitness, it is perceived as a stressful experience manifested by the appearance of displacement-like behavior and accompanied by temporal costs in the form of sensitivity to acute stressors. This is the first example in flies of a situation in which deprivation of sexual reward is not simply a lack of reward but is perceived as a stressful experience.

In search for mechanisms that could explain the behavioral and physiological responses to deprivation of sexual reward we examined four possible directions: (1) changes in gene expression within discrete neuronal populations (2) energetic costs or modulation of metabolic pathways (3) activation of cytoprotective stress response (4) disinhibition of NPF target neurons. Although Gendron et al. (2014^75^) showed that exposure to female, as opposed to male, pheromones lowers the levels of TAG in males, we did not detect metabolic changes that could explain sensitivity to starvation or oxidative stress, and therefore ruled out energetic cost as the cause for sensitivity to starvation stress. Still, the metabolome highlighted some interesting metabolites, such as 5-Aminolevulinic acid, which is enriched in rejected flies. This may indicate a reduction in Hem synthesis and consequently an elevation in protoporphyrin, and a possible reduction in heme oxigenase (HO). Although there is some evidence that protoporphyrin can act as an antioxidant^121,122^ it mainly functions as a pro-oxidative^123,124^. HO is a rate-limiting enzyme that degrades heme into biliverdin, carbon monoxide (CO), and iron^125^. In *Drosophila, ho* gene is expressed in different brain tissues including the optic lobe, the central brain, and glial cells, and plays an important role in cell survival and protection against paraquat-induced oxidative stress^126^. This is consistent with rejected males experiencing oxidative stress and are thus more sensitive to paraquat. However, this does not clarify why in addition, rejected males were more sensitive to starvation.

We previously generated a transcriptomic map of the fly brain, demonstrating that different neuronal populations possess distinct transcriptional programs^60^, and extended this approach to identify changes in gene expression within selected neuronal populations following social and sexual experience. An example of the strength of the approach is the downregulation of Ubiquitin conjugating enzyme (UBC7) in NPFR neurons in response to rejection. Though a null mutation of UBC7 that eliminates its expression in all tissue disrupts the ability of male flies to court^89^, its knockdown in NPFR neurons leads to the opposite phenotype, including enhanced motivation to court and increased social arousal^60^, providing a functional explanation for its downregulation in response to rejection.

Though the INTACT technology provides valuable information that broadens our understanding of the molecular signature of neurons under different conditions, it is challenging to bridge the gap between DEGs and their functional relevance. While imaging NPFR^TK^ neurons, we noticed that their expression did not match the known tk expression^41,51,91,127,128^. This could be attributed to the high resolution of INTACT, which enabled us to identify a minor but significant elevation of tk transcript in NPFR cells whereas protein/peptide could not be identified by immunolabeling. Alternatively, this could result from a tk promoter activity at a very specific period in fly development, facilitating the expression of LexAop-FlpL in these cells. Another explanation is that only tk transcript is upregulated, but no TK peptides are generated.

One of the limitations of the INTACT method that makes it difficult to interpret the genomic results, is that it uses genetic drivers to mark certain neuronal populations. The drivers are based on the expression pattern of marker genes such as NPF or NPFR, resulting in the isolation of cells that express this marker and are composed of smaller subsets of neurons with composite molecular signatures. By tagging neurons using this approach, we extract all the cells that express a certain marker gene, inevitably pooling together very different cells in terms of function and location inside the brain. A good example for this is the NPFR driver, which marks a population of 100 neurons, some of which also express NPF^120^, while other express Dh44, TK or are dopaminergic^59^. A way to overcome this limitation is profiling smaller numbers of neurons using an improved version of the INTACT called TAPIN^129^ and new technologies that allow mapping of the expression of single RNA molecules in intact brain tissue, such as expansion sequencing^130^.

We revealed an anatomical association between reward and stress systems in the form of receptors for NPF on Dh44 neurons, resembling the known interplay between NPY and CRF neurons in mammals^29^. In addition, we documented a translocation of FoxO to the cytoplasm of Dh44 neurons in response to deprivation of sexual reward followed by starvation. Although this could represent the activation of a cellular stress response that could limit the ability of flies to resist other types of stressors, as has been shown in C.elegans^109^, we did not find a functional connection between rejection, FoxO translocation in Dh44 neurons and increased sensitivity to starvation.

After ruling out metabolic changes and brain cytoprotective stress response as possible explanations for the responses of male flies to deprivation of sexual responses, we demonstrated that sensitivity to starvation stress is caused by disinhibition of NPF target neurons, mimicked in our experiments by optogenetic activation of NPFR neurons as well as KD of *npfr* transcript in NPFR neurons. This response is mediated through a dynamin independent signaling mechanism, possibly bulk endocytosis or neuropeptide release, which does not involve synaptic vesicle reuptake. Interestingly, single males were not affected by NPFR KD. This can be attributed to their lack of sexual and social interaction. This can be also seen in their distinct internal state reflected by a unique transcriptome and metabolome expression. Furthermore, the activation of only ~22-26 NPFR^TK^ neurons proved sufficient to induce both increased sensitivity to starvation and some of the behavioral phenotypes observed in rejected males. The identified NPFR sub-population can be divided into smaller known subpopulations: six of these neurons express Dh44, and two pairs of NPFR^TK^ neurons, L1-l and P1, colocalize with NPF expressing neurons (Fig. 6F). Both of these NPFR-NPF neuron pairs were previously indicated as not related to wakefulness promotion, and presumably not involved in reward-seeking behavior^76,120^. Thus, it is likely that L1-l, P1 NPFR-NPF neurons do not facilitate increased arousal and might not be affected by sexual reward deprivation. Indeed, we showed that the activation of these neurons did not affect either resistance to starvation or group behavior in the flybowl. While CRF neurons in mammals mediate reward seeking behaviors and responses to social stress^21,35–37,131–136^ and the fly Dh44 neurons are known to facilitate aggressive behaviors^69^, we did not find evidence to support their role in regulating responses to social stress. Therefore, the activity of NPF\NPFR circuit alone is sufficient to facilitate a stress-like response caused by deprivation of a reward and increase subsequent sensitivity to acute stressors (Fig. 6G). Further dissection of the NPFR and NPFR^TK^ neuronal subpopulations is needed to better understand the role of each cell type in response to sexual reward deprivation.

This is the first documentation of a frustration-like stress response in *Drosophila,* which is caused by innate expectation of a natural reward that is not achieved. Sexual deprivation, therefore, can be used as a tool to study the interplay between natural reward omission, reward seeking behaviors such as ethanol consumption, stress, and addiction. Further investigation of cell type-specific mechanisms that facilitate frustration-like response to reward omission will shed more light on the complex mechanism by which the brain processes motivation and reward.

## Methods

### Fly lines and culture

*Drosophila melanogaster* WT Canton S flies were kept at 25C°, ~50% humidity, light/dark of 12:12 hours, and maintained on cornmeal, yeast, molasses, and agar medium. Most fly lines were back-crossed to a Canton S background. UAS *UAS.unc84-2XGFP* and UAS mCD8 GFP were obtained from HHMI Janelia Research Campus. For INTACT, UAS.*unc84*-2XGFP transgenic flies were crossed with NPFR-GAL4 flies. NPFR-Gal4, UAS-NPFR RNAi, NPF-GAL4 flies were a gift from the Truman lab (HHMI Janelia Campus). UAS<dsFRT>cs-Chrimson-mVenus in attp2, and LexAop-FLPL flies were a gift from the Heberlein lab (HHMI Janelia Campus). UAS-dFoxO-TM flies were a kind gift from the Tatar lab (Brown University). The following lines were ordered from the indicated fly centers: TK-LexA (Bloomington #54080), UAS-tk-HA (ORF F000997), UAS-tk RNAi (VDRC #103662), Dh44-Gal4 (VDRC #207474).

### RNA extractions from different neuronal cell types (INTACT)

Cell type specific labeled nuclei were isolated using the INTACT method (Isolation of Nuclei Tagged in A specific Cell Type technique) as previously described^87^. This method was slightly modified as follows: about 100 adult male flies collected from 3-4 days F1 generation of NPFR GAL4_driver X UAS_*unc84*_2XGFP_ reporter were subjected to a courtship assay, at the end of which they were anesthetized by CO2 and their heads were separated on ice using a scalpel. 9ml of homogenization buffer (20mM β-Glycerophosphate pH7, 200mM NaCl, 2mM EDTA, 0.5% NP40 supplemented with RNAase inhibitor,10mg/ml t-RNA, 50mg/ml ultrapure BSA, 0.5mM Spermidine, 0.15mM Spermine and 140ul of carboxyl Dynabeads −270 (Invitrogene: 14305D) was added to each sample. The heads were filtered on ice by a series of mechanical grinding steps followed by filtering the homogenate using a 10um Partek filter assembly (Partek: 0400422314). After removing the carboxyl-coated Dynabeads using a magnet, the homogenate was filtered using a 1um pluriSelect filter (pluriSelect: 435000103). The liquid phase was carefully placed on a 40% optiprep cushion layer and centrifuged in a 4°C centrifuge for 30min at ~2300Xg. The homogenate/Optiprep interface was incubated with anti-GFP antibody (Invitrogen: G10362) and protein G Dynabeads (Invitrogen: 100-03D) for 40 minutes at 4°C. Beads were then washed once in NUN buffer (20mM β-Glycerophosphate pH7, 300mM NaCl, 1M Urea, 0.5% NP40, 2mM EDTA, 0.5mM Spermidine, 0.15mM Spermine, 1mM DTT, 1X Complete protease inhibitor, 0.075mg/ml Yeast torula RNA, 0.05Units/μl Superasin). Bead-bound nuclei were separated using a magnet stand and resuspended in 100μl of RNA extraction buffer (Picopure kit, Invitrogen # KIT0204), and RNA was extracted using the standard protocol.

### RNA-seq library preparation and sequencing

The NuGEN RNAseq v2 (7102-32) kit was used to prepare cDNA from the INTACT purified RNA, followed by library preparation using the SPIA – NuGEN Encore Rapid DR prep kit. Samples were sequenced on an Illumina HiSeq using single-end 60 base pair reads.

### Determining gene expression levels from RNA-seq

Reads were trimmed using cutadapt^137^ and mapped to *Drosophila melanogaster* (BDGP6) genome using STAR^138^ v2.4.2a (with EndToEnd option and outFilterMismatchNoverLmax was set to 0.04). Counting proceeded over genes annotated in Ensembl release 31, using htseq-count^139^ (intersection-strict mode). Reads overlapping exons in each gene were counted using featureCounts^140^, and these counts were used as input into DESeq2^141^. DeSeq2 function rlog(blind=FALSE) was used to calculate normalized counts with a regularized log transformation. The DESeq() and results() functions were used to calculate gene expression differences between pairs of cell types.

### Behavioral assays: 1. Courtship suppression recording

WT males and females were collected on CO2 3-4 days before the recording. Males were kept in groups of 25 per vial. To generate mated females for the experiment, virgin females were introduced to males ~16 hours before the experiment. All flies were kept in the incubator at 25°C, ~50% humidity, and light/dark of 12:12 hours. Prior to the conditioning, the mated females were separated from the males on CO2 on the morning of the recording. During the recording, the temperature was kept at ~25°C, and humidity ~55%. Since the extent of courtship display is shaped by circadian rhythmicity, where male flies depict the highest courtship activity closest to the onset of light, and their general activity declines towards noon, the first session started right after the onset of light, and the other two sessions took place in the afternoon. Virgin male flies were exposed to either mated or virgin female for three one-hour sessions, and their behavior was recorded using a Point Grey Firefly camera and analyzed in detail during the first 10 minutes of each interaction. At the end of each session, female flies were removed, and the males that experienced rejection were kept isolated in narrow glass vials for one hour. At the end of the rest hour, males were returned to their original location in the courtship arena for the recording. In order to compare the courtship behavior of rejected and naive males, virgin males from the naive cohort were replaced at the beginning of each session. Different aspects of courtship behavior were analyzed manually using “Lifesong” software.

### 2.Courtship conditioning for INTACT

Males and females were collected within 2 h of eclosion on CO2, 3-4 days before courtship conditioning. Males were collected into narrow glass vials (VWR culture glass tubes 10X75mm) containing food and kept single housed until the conditioning. To generate mated females for the experiment, mature males were added to the females ~16 h before the experiment. All flies were kept in the incubator at 25°C, ~50% humidity, and light/dark of 12:12 hours. The mated females were separated from the males on the morning of the conditioning. During the conditioning, the temperature was kept at about 25°C, and humidity ~55%.

**Generation of rejected males**: Individual males were placed with mated females for 3 one-h conditioning trials (separated by 1-h rests) a day for two consecutive days. Females were removed after each trial. **Generating mated males:** To generate the “mated-grouped” cohort, individual males were housed with virgin females for 3 one-h conditioning trials (separated by 1-h rests) a day for two consecutive days. Females were removed after each trial. **Single males:** Virgin males were collected within 2 h of eclosion and kept separately in small food vials during the entire trial. Gentle handling was performed parallel to rejected and mated males’ conditioning sessions.

### 3.FlyBowl

FlyBowl experiments were conducted as described in Bentzur et. al^119^. In brief: groups of 10 male flies, which were socially raised in groups of 10 for 3-4 days, were placed in FlyBowl arenas, and their behavior was recorded at 30 fps for 15 min and tracked using Ctrax^142^. Automatic behavior classifiers and Per-frame features were computed by JABBA^143^ tracking system. Data of all behavioral features were normalized to percentage of difference from the average of each experiment for visualization. Details about the different features are found in Figure S4.

### Optogenetics Activation of NPFR, Dh44, and NPFR^TK^ neurons

Light-induced activation of red-shifted Channel Rhodopsin UAS-CsCrimson was achieved by placing glass fly vials containing one fly each over red LEDs (40 Hz, 650nm, 0.6 lm @20mA). Activation protocol consisted of 3×5 min-long activation periods spaced by 1 h and 55 min resting intervals for 2 consecutive days.

### Neuronal activation combined with inhibition of synaptic vesicle release

Flies expressing Cs-Chrimson and UAS-Shibire^ts^ in NPFR neurons were subjected to one of four conditions for two days: (1) Three 5-min-long optogenetic activations spaced by 1 h and 55 min resting intervals (under constant dark) at constant 18-20°C served as a positive control. (2) Three 10-min-long sessions at 28-29°C under constant dark followed by 5 min long optogenetic activations spaced by 1 h and 45 min resting intervals at 18-20°C, also under constant dark. (3) Three 15-min-long sessions at 28-29°C under constant dark, spaced by 1 h and 45 min at 18-20°C, also under constant dark, served as synaptic release block control. (4) Flies kept at constant 18-20°C and constant dark served as a negative control. After the last activation, flies were transferred into glass vials containing 1% agarose.

### Immunostaining

Whole-mount brains were fixed for 20 min in 4% paraformaldehyde (PFA) or over-night in 1.7% PFA. Preparations were blocked for 1h at 4°C with gentle agitation in 0.5% BSA, 0.3% Triton in PBS. The following primary antibodies were used: Rabbit anti GFP (LifeTech 1:500), the neuropile-specific antibody NC82, (1:50, The Jackson Laboratory), mouse anti GFP (1:100, Roche), Rabbit anti Dh44 (0.6:100), rabbit anti dILp2 (1:100, a kind gift from Takashi Nishimura lab), rabbit anti FoxO (1:100. A kind gift from Pierre Leopold lab), rabbit anti tk (1:1000 a Kind gift from Wei Song) were incubated overnight at 4°C. Secondary antibodies, goat anti mouse-Alexa488 (1:200-1:100), goat anti rabbit-Alexa568 (1:200-1:100), goat anti mouse-Alexa568 (1:1000) and goat anti rabbit-Alexa488 (1:1000) were incubated for 2hr at 4°C. DAPI (1:20). The stained samples were mounted with SlowFade^TM^ Gold antifade reagent (Thermo Fisher Scientific) and visualized using a Leica SP8 confocal microscope.

### Stress tests

Survival was measured during: <u>(i) Starvation</u>: After the last bout of conditioning, males were transferred to glass vials containing 1% agarose. Males were kept in isolation, and the number of living flies was recorded every 2-3 h.

### (ii)Oxidative stress

To induce oxidative stress, male flies were single-housed in glass vials containing standard food supplemented with 20mM paraquat (856177,Sigma-Aldrich).

### Longevity

After the last bout of conditioning, males were transferred to vials containing food. Males were kept in isolation and the number of living flies was recorded every day. Flies were transferred to new vials twice a week. Log rank or Renyi-type test (REF) with FDR correction were performed.

### TAG, Glucose levels evaluation

TAG levels were assessed as described^75^ with modifications: After courtship conditioning, experimental males were divided into groups of 5 and were homogenized together in 100 μl NP40 substitute assay reagent from Triglyceride colorimetric assay kit 10010303 (Cayman JM-K622-100). Homogenate was centrifuged at 10,000 x g for 10 min at 4°C, and the supernatant was collected. Triglyceride enzyme mixture (10010511) was used to hydrolyze the triglycerides and subsequently measure glycerol by a coupled enzymatic reaction. TAG concentrations were determined by the absorbance at 540nm and estimated by a known triglyceride standard. The absorbance was measured using SynergyH1 Hybrid Multi-Mode microplate Reader.

Body and hemolymph glucose were extracted as described^144^. Briefly (with modifications):

**Whole bodies:** After courtship conditioning, 5 males were placed in each sample tube and were weighed using Fisher scientific ALF104 analytical balance scale. Then, flies were homogenized in 100 ml cold PBS on ice. Supernatant was heated for 10 min at 70°C, then centrifuged for 3 min at maximum speed at 4°C. Supernatant was collected and transferred to a new 1.5 ml tube. **Hemolymph:** After courtship conditioning, males were sedated on ice and carefully punctured in the thorax using sharpen stainless steel tweezers. 40 punctured flies were placed in each 0.5 ml microfuge tube with a hole at the bottom made by a 25G needle. The 0.5 microfuge tube was then placed in a 1.5 ml tube and centrifuged at 5000rpm for 10 min at 4°C. Hemolymph was collected, and samples were heated for 5 min at 70°C. Glucose was measured using High sensitivity Glucose Assay kit (MAK181 Sigma-Aldrich). Glucose concentration is determined by coupled enzyme assay, which results in fluorometric (lex = 535/lem = 587nm) products and was assessed using SynergyH1.

### Metabolites extraction

After courtship conditioning, males were flash-frozen and decapitated using a microscalpel. For each samples, 5 heads were transferred into soft tissue homogenizing CK 14 tubes containing 1.4 mm ceramic beads (Bertin corp.) prefilled with 600 ul of cold (−20 °C) metabolite extraction solvent containing interanl standards (Methanol:Acetonitrile:H2O::50:30:20) and kept on ice. Samples were homogenized using Precellys 24 tissue homogenizer (Bertin Technologies) cooled to 4°C (3 × 30s at 6000 rpm, with a 30 s gap between each cycle). Homogenized extracts were centrifuged in the Precellys tubes at 18,000 g for 10 min at 4 °C. The supernatants were transferred to glass HPLC vials and kept at −75 °C prior to LC-MS analysis.

### LC-MS metabolomic analysis

LC-MS analysis was conducted as described^145^. Briefly, Dionex Ultimate ultra-high-performance liquid chromatography (UPLC) system coupled to Orbitrap Q-Exactive Mass Spectrometer (Thermo Fisher Scientific) was used. Resolution was set to 35,000 at 200 mass/charge ratio (m/z) with electrospray ionization and polarity switching mode to enable both positive and negative ions across a mass range of 67-1000 m/z. UPLC setup consisted ZIC-pHILIC column (SeQuant; 150 mm × 2.1 mm, 5 μm; Merck). 5 μl of cells extracts were injected and the compounds were separated using a mobile phase gradient of 15 min, starting at 20% aqueous (20mM ammonium carbonate adjusted to pH 9.2 with 0.1% of 25% ammonium hydroxide): 80% organic (acetonitrile) and terminated with 20% acetonitrile. Flow rate and column temperature were maintained at 0.2ml/min and 45 °C, respectively, for a total run time of 27 min. All metabolites were detected using mass accuracy below 5 ppm. Thermo Xcalibur 4.1 was used for data acquisition. The peak areas of different metabolites were determined using Thermo TraceFinder^TM^ 4.1 software, where metabolites were identified by the exact mass of the singly charged ion and by known retention time, using an in-house MS library built by running commercial standards for all detected metabolites. Each identified metabolite intensity was normalized to ug protein. Metabolite-Auto Plotter^146^ was used for data visualization during data processing.

### FoxO in cyto\nuc

Dissected stained samples of male’s brains were visualized using a Leica SP8 confocal microscope, with an X60 lens. Images of cells containing both immunostaining to FoxO and DAPI were analyzed using CellProfiles 3.1.9. Images were acquired with different acquisition settings because of the very large dynamic range difference among the samples. Cell cytoplasm intensity was calculated as (whole-cell intensity) – (nucleus intensity). To quantify the differences among samples, intensity was measured in each cell in the nucleus (nuc) and cytoplasm (cyto), and the ratio was then calculated as the relation between the cytoplasm and nucleus in order to normalize. This ratio was compared among samples. The cyto\nuc portion of rejected, mated, and single males was compared and analyzed using Kruskal-Wallis and post-hoc Dunn’s test.

### Statistical analysis

Data of each behavioral feature per experiment were tested for normality, and consequently, normally distributed data were tested by student’s t-test, one-way ANOVA followed by Tukey’s post-hoc. Non-parametric data were tested by Mann-Whitney or Kruskal-Wallis tests followed by Dunn’s or Friedman’s post-hoc tests. FDR correction for multiple comparisons was performed for all Flybowl experiments features. Statistical overrepresentation was generated using PANTHER^147,148^ (http://pantherdb.org/citePanther.jsp). Kmeans clustering method was performed (k= 3) to generate a heatmap of differentially expressed genes in NPFR neurons. Starvation resistance and longevity experiments were tested by Log-rank or Renyi-type test^149^ using R package version 3.2-11. FDR correction for multiple comparisons was performed for experiments with more than two experimental groups.

## Acknowledgments

We thank all members of the Shohat-Ophir lab for fruitful discussions and technical support. We would also like to thank Jennifer I. C. Benichou for statistical consultation. This work was supported by the Israel Science Foundation Grant 384/14 and Israel Science Foundation Grant 174/19.

---

**Figure S1.**
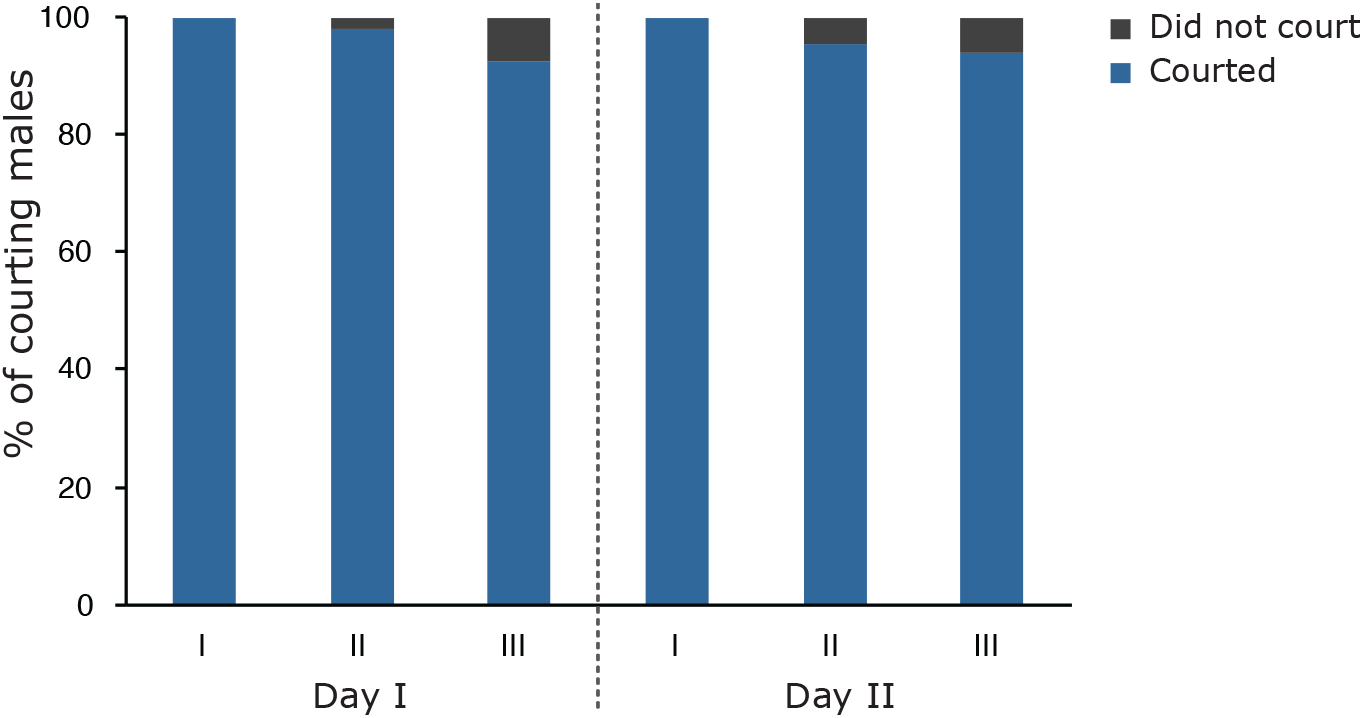
Repeated rejection does not lower the percentage of males who are still motivated to court. Naive males were repeatedly introduced to sexually non-receptive females over two consecutive days. The number of courting males was documented for each session. Males that did not initiate courtship, or that succeeded to copulate were excluded from further analysis. Total n of males for day I= 97 first session (97 courted), 97 second session (95 courted), 95 third session (88 courted), n for day II= 86 first session (86 courted), 86 second session (82 courted), 82 third session (77 courted). Bar graph represents % of courting males in each session.

**Figure S2.**
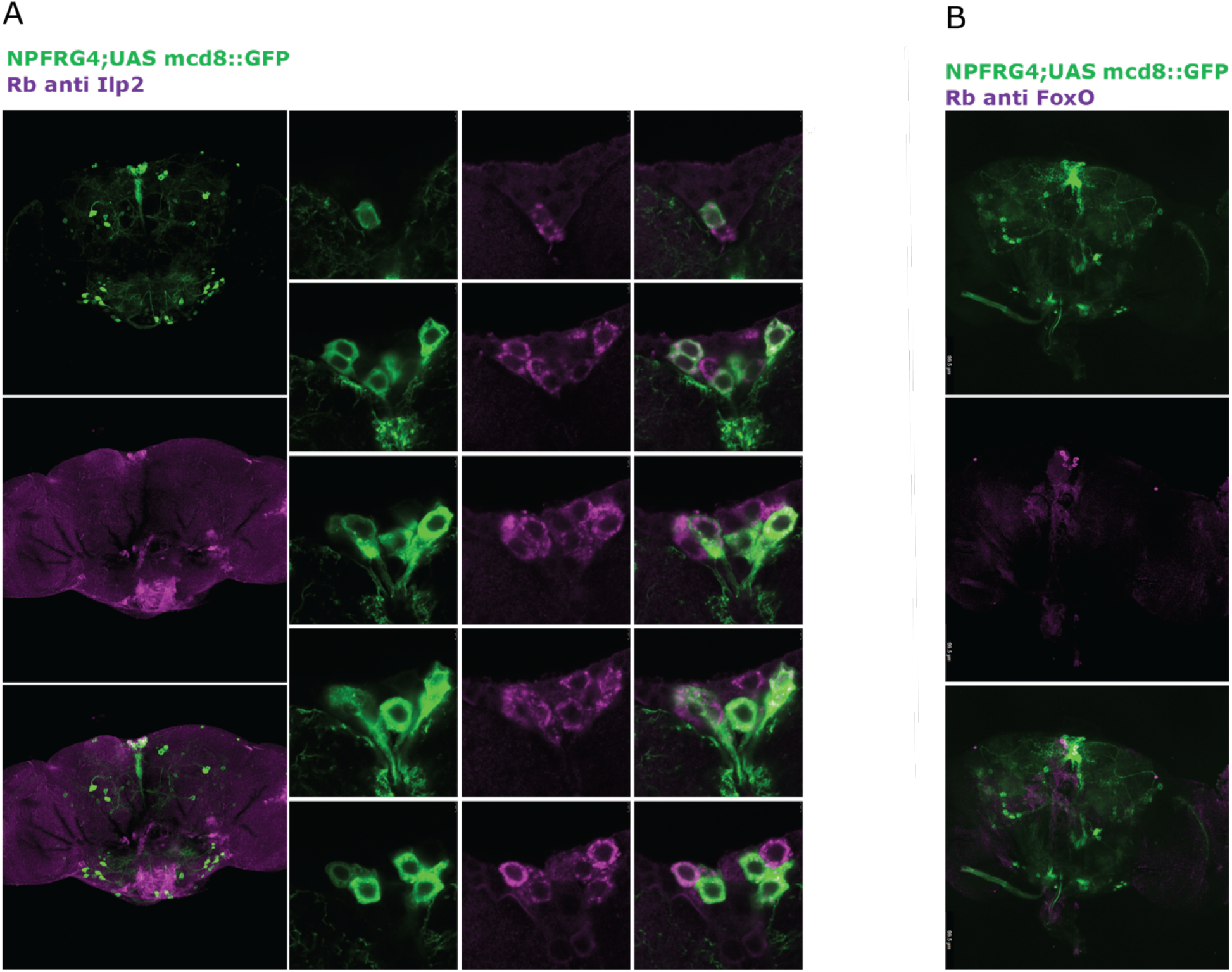
NPFR neurons colocalize with Ilp2 and FoxO expressing neurons. **A**. Immunostaining of NPFRG4; mcd8::GFP neurons using antibodies to GFP (green) and endogenous Ilp2 (magenta). On the left: whole brains. On the right: cells in the PI. **B**. Immunostaining of NPFRG4; mcd8::GFP neurons using antibodies to GFP (green) and endogenous FoxO (magenta).

**Figure S3.**
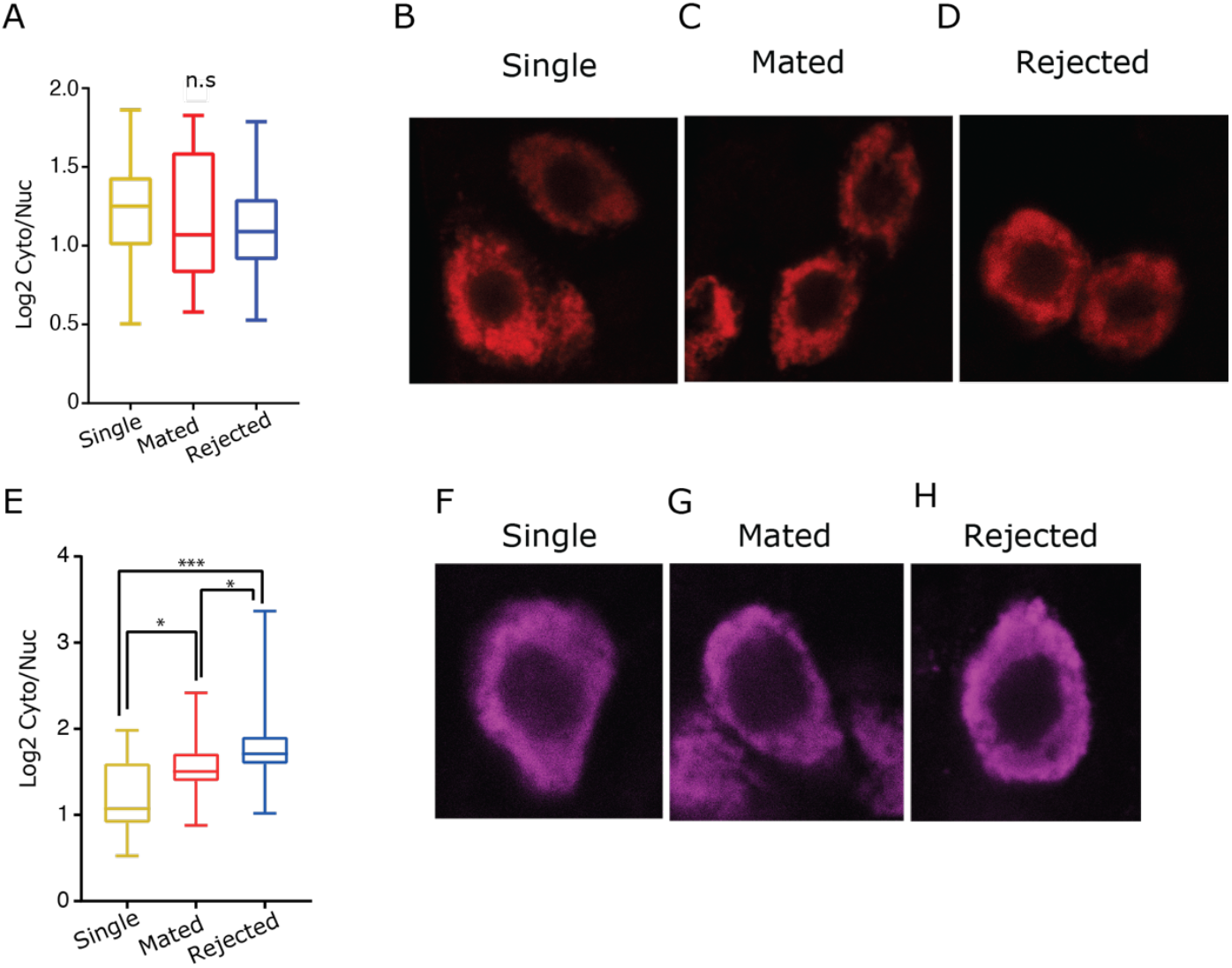
Endogenous FoxO localizes to the cytoplasm in starved rejected males. **A.** The ratio of cytoplasmic/nuclear localized FoxO in rejected males (blue, n=33 cells (7 brains)) compared to mated (red, n=18 cells (5 brains)), and single males (yellow, n=29 cells (5 brains). *p*>0.05 Kruskal-Wallis with Friedman post-hoc, No significant differences were observed. **B-D.** FoxO positive cells (red) of single (**B**), mated (**C**), and rejected (**D**) males. **E.** The ratio of cytoplasmic/nuclear localized FoxO in rejected males (blue, n=34 cells (6 brains)) compared to mated (red, n=30 cells (6 brains)) and single (yellow, n=33 cells (6 brains)) males, all of which were starved for 20 h. Rejected vs mated, mated vs single, \**p*<0.05, rejected vs single, \*\*\*\**p*<0.0001. Kruskal-Wallis with Friedman post-hoc was performed. **F-H**. FoxO positive cells (magenta) of starved single (**F**), mated (**G**), and rejected (**H**) males.

**Figure S4.**
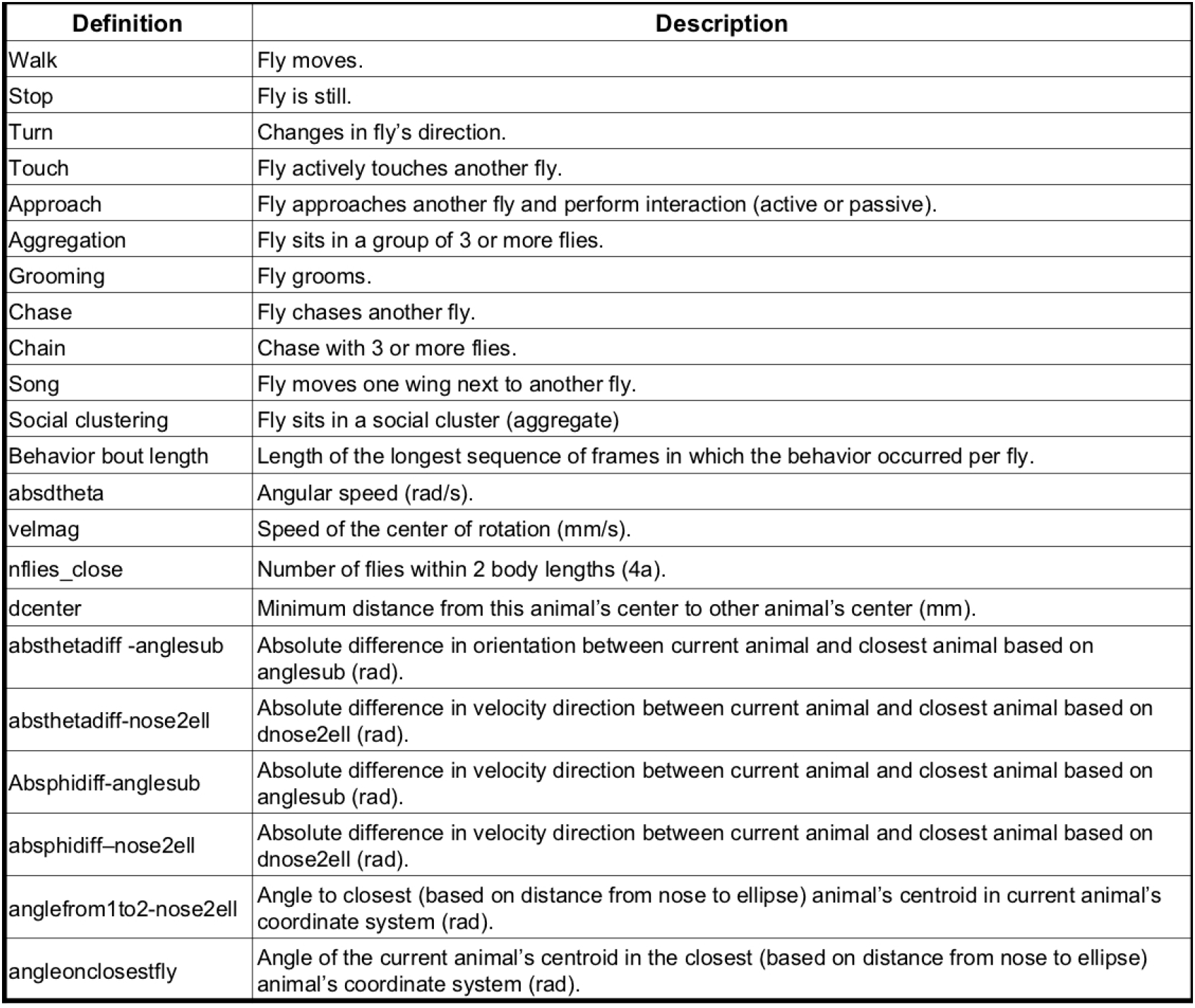
List of behavioral features presented in Figures 5, 6 and S6.

**Figure S5.**
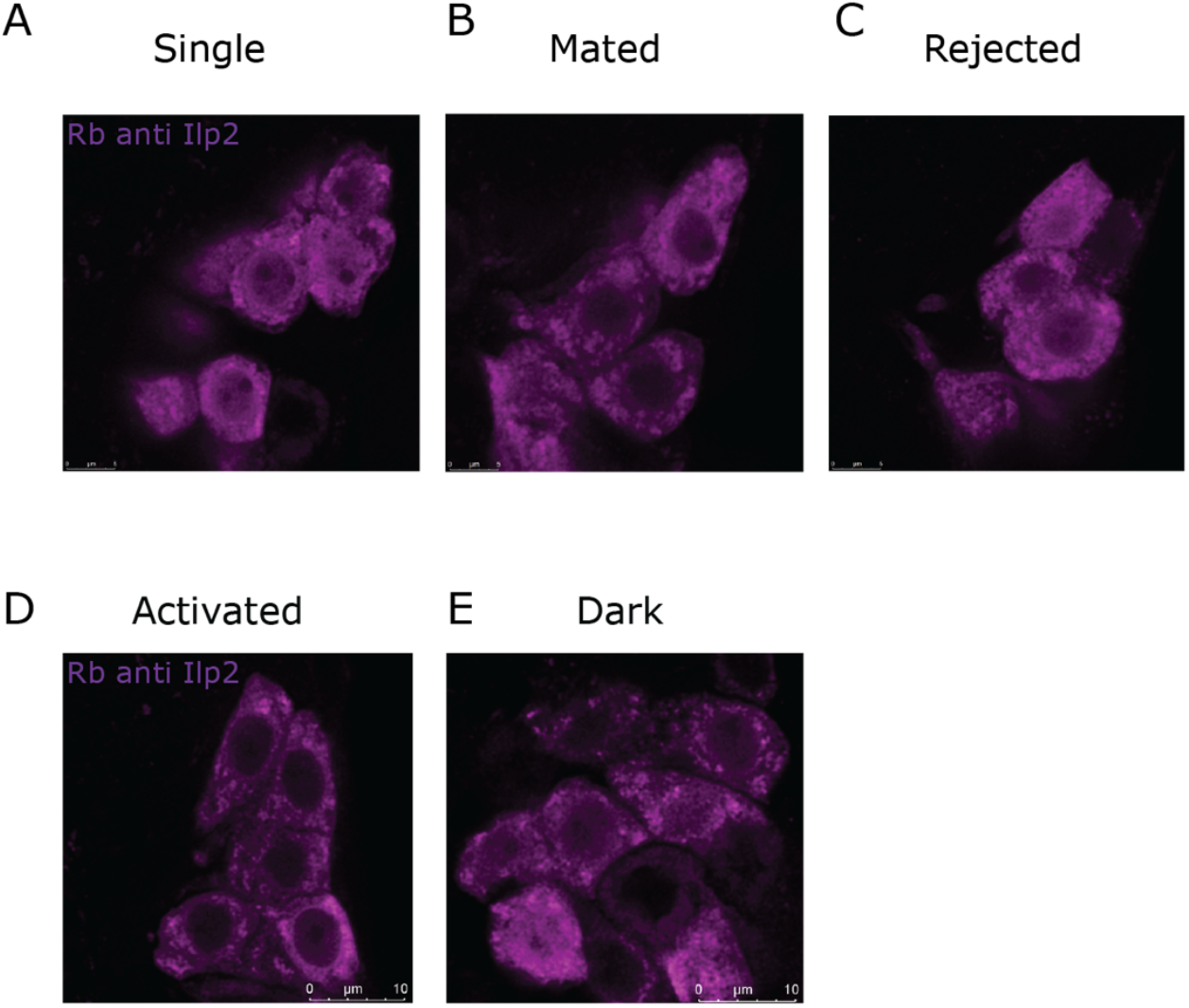
Activation of NPFR neurons does not affect insulin peptide 2 (Ilp2) abundance in IPCs. **A.-C.** Immunostaining of male brains using antibodies to endogenous Ilp2 following courtship conditioning. Fluorescent microscope imaging of IPCs in single (A), mated (B), and rejected (C) males. **D,E.** NPFR neurons were activated in NPFR G4/+;UAS-csChrimson/+ male flies by exposing them to red light three times a day for two days (D). Control flies were not exposed to light (E). Brains of both groups were stained with antibodies to endogenous Ilp2 and imaged.

**Figure S6.**
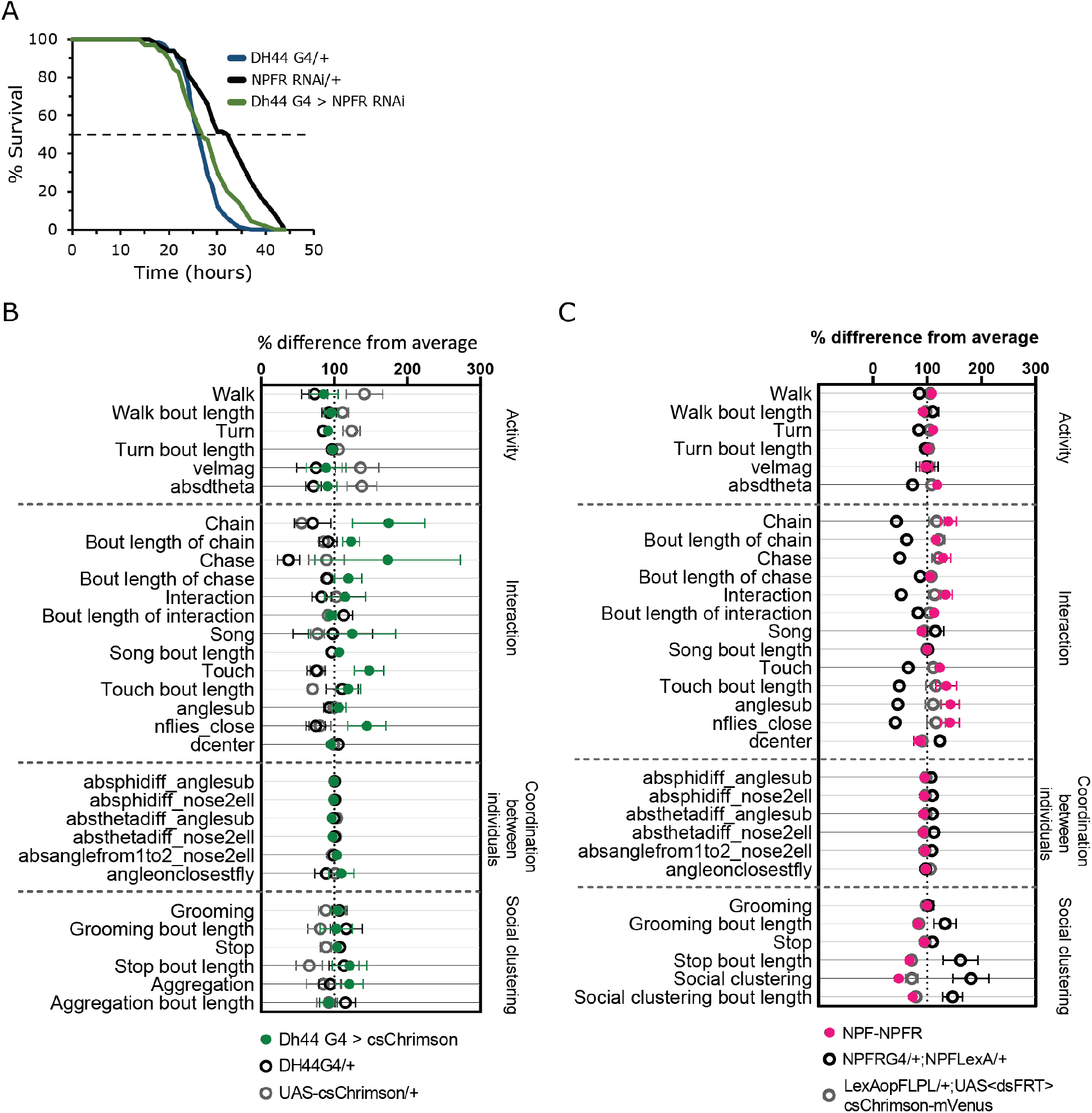
Disinhibition of DH44 or NPFR-NPF neurons did not affect male group behavior. **A.** Starvation resistance assayed on Dh44 G4/NPFR RNAi (green, n=70) flies and their genetic controls Dh44 G4/+ (blue, n=66) and NPFR RNAi/+ (black, n=64). No significant difference was observed among experimental flies and the controls, *p*>0.05. Pairwise log-rank test with FDR correction for multiple comparisons was performed. **B.** % difference average scatter plot of behaviors observed and scored in the FlyBowl performed by DH44 G4/+;UAS-csChrimson/+ males (green, n=8) and their genetic controls Dh44 G4/+, UAS-csChrimson/+ (black, n=7 and grey, n=8, respectively). \**p*<0.05. Kruskal-Wallis and pairwise log-rank with FDR correction for multiple comparisons were performed. **C.** % difference average scatter plot of behaviors observed and scored in the FlyBowl performed by NPFR-NPF males (pink, n=22) and their genetic controls NPFR G4/+;NPF LexA/+ (grey, n=22), LexAop-FLPL/+;UAS<dsFRT>cs-Chrimson-mVenus/+, (black, n=21). *p*>0.05 Kruskal-Wallis. ANOVA or Kruskal-Wallis with post-hoc Tukey’s or Dunn’s test, and FDR correction for multiple comparisons were performed.

**Figure S7.**
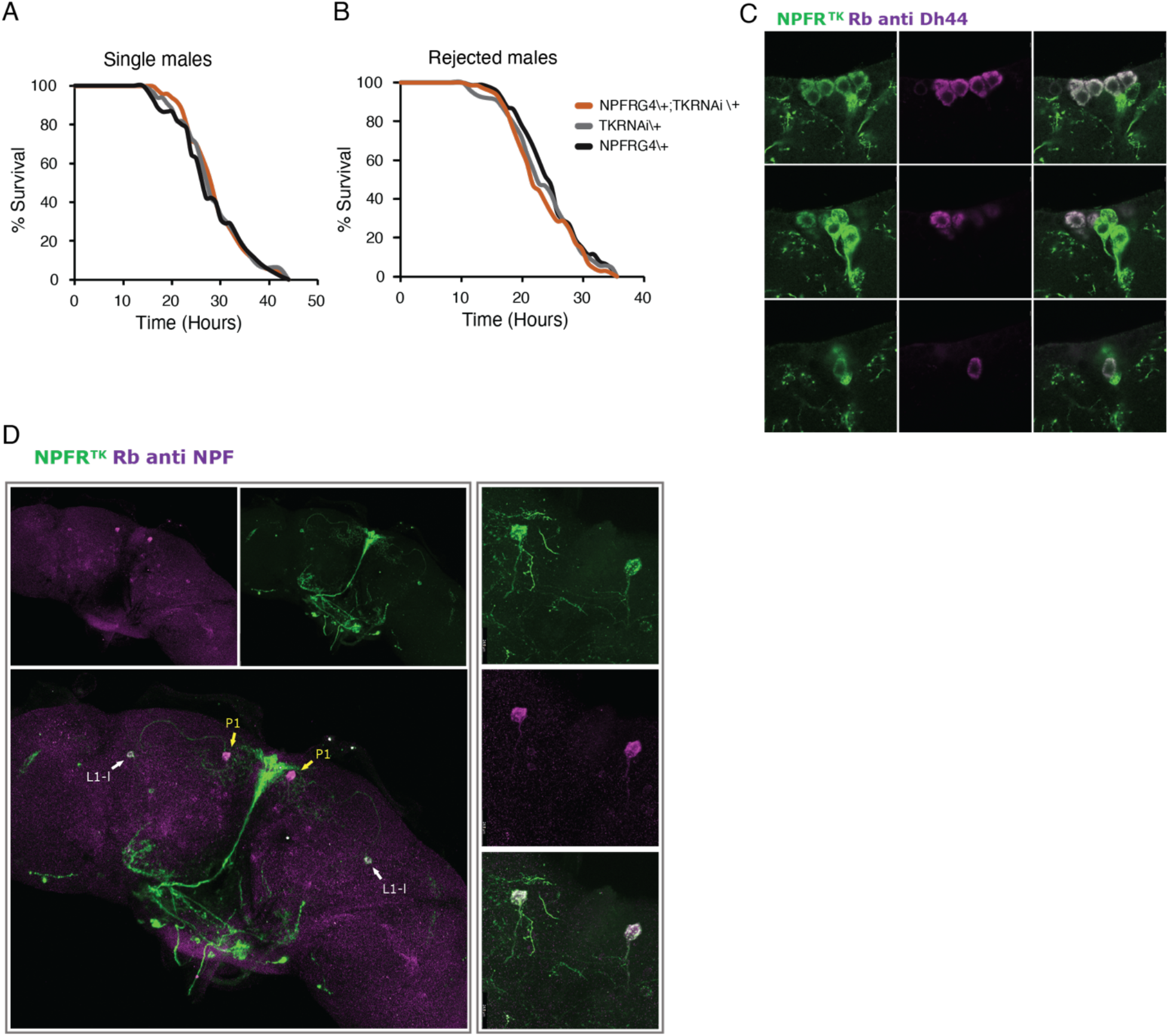
TK expression in NPFR neurons does not affect resistance to starvation. **A.** KD of tk in NPFR neurons of naïve males using NPFRG4/+;tkRNAi/+. Experimental single housed NPFR G4/+;TK RNAi/+ (orange, n=50) flies did not exhibit different resistance to starvation compared to genetic controls: TK RNAi/+ (grey, n=33) and NPFR G4/+ (black, n=45). **B.** NPFRG4/+;tkRNAi/+ males and their genetic controls were subjected to rejection and their resistance to starvation was assayed. No significant difference in resistance to starvation in NPFR G4;tk RNAi flies (orange, n=79) compared to genetic controls (grey, n=90 and black, n=70) was observed. Pairwise log-rank test with FDR correction for multiple comparisons was performed for A,B. **C.** Six NPFR^TK^ (green) neurons colocalize with DH44 (magenta, endogenous Dh44 expression). **D. Right**: colocalization of NPFR^TK^ neurons (green) and NPF+ neurons (magenta, endogenous NPF expression), indicated by arrows. White arrows indicate L1-l neurons, yellow arrows indicate P1 neurons. **Left:** A closeup to two NPFR^TK^ NPF+ neurons (P1).

## Supplementary Tables

**Table S1**: List of differentially expressed genes shown in figure 2C. Genes were clustered using Kmeans clustering method (k= 3).

**Table S2**: List of differentially expressed genes between rejected, mated and single cohorts.

**Table S3**: LC-MS metabolite results. Compound averaged peak area/total measurable ions results are presented for mated (n= 16), rejected (n= 17), and single (n= 17) cohorts.

## References

1. O’Connell, L. A. & Hofmann, H. A. The Vertebrate mesolimbic reward system and social behavior network: A comparative synthesis. Journal of Comparative Neurology vol. 519 3599–3639 (2011).

2. Kim, S. M., Su, C.-Y. & Wang, J. W. Neuromodulation of Innate Behaviors in Drosophila. Annu. Rev. Neurosci. 40, 327–348 (2017).

3. Shemesh, Y., Sztainberg, Y., Forkosh, O., Shlapobersky, T., Chen, A. & Schneidman, E. High-order social interactions in groups of mice. Elife 2013, (2013).

4. Aureli, F. & Schino, G. Social complexity from within: how individuals experience the structure and organization of their groups. Behav. Ecol. Sociobiol. 2019 731 73, 1-13 (2019).

5. Forkosh, O., Karamihalev, S., Roeh, S., Alon, U., Anpilov, S., Touma, C., Nussbaumer, M., Flachskamm, C., Kaplick, P. M., Shemesh, Y. & Chen, A. Identity domains capture individual differences from across the behavioral repertoire. Nat. Neurosci. 2019 2212 22, 2023–2028 (2019).

6. Hobson, E. A., Ferdinand, V., Kolchinsky, A. & Garland, J. Rethinking animal social complexity measures with the help of complex systems concepts. Anim. Behav. 155, 287–296 (2019).

7. Kravitz, E. A. & Huber, R. Aggression in invertebrates. Current Opinion in Neurobiology vol. 13 736–743 (2003).

8. Anderson, D. J. Circuit modules linking internal states and social behaviour in flies and mice. Nat. Rev. Neurosci. 17, 692–704 (2016).

9. Koob, G. & Kreek, M. J. Stress, dysregulation of drug reward pathways, and the transition to drug dependence. American Journal of Psychiatry vol. 164 1149–1159 (2007).

10. Papini, M. R. & Dudley, R. T. Consequences of surprising reward omissions. Rev. Gen. Psychol. 1, 175–197 (1997).

11. Do-Monte, F. H., Minier-Toribio, A., Quiñones-Laracuente, K., Medina-Colón, E. M. & Quirk, G. J. Thalamic Regulation of Sucrose Seeking during Unexpected Reward Omission. Neuron 94, 388–400.e4 (2017).

12. Zimmerman, P. H. & Koene, P. The effect of frustrative nonreward on vocalisations and behaviour in the laying hen, Gallus gallus domesticus. Behav. Processes 44, 73–79 (1998).

13. Burokas, A., Gutiérrez-Cuesta, J., Martín-García, E. & Maldonado, R. Operant model of frustrated expected reward in mice. Addict. Biol. 17, 770–782 (2012).

14. Duncan, I. J. H. & Wood-Gush, D. G. M. Frustration and aggression in the domestic fowl. Anim. Behav. 19, 500–504 (1971).

15. Dantzer, R., Arnone, M. & Mormede, P. Effects of frustration on behaviour and plasma corticosteroid levels in pigs. Physiol. Behav. 24, 1–4 (1980).

16. De Almeida, R. M. M. & Miczek, K. A. Aggression escalated by social instigation or by discontinuation of reinforcement (‘Frustration’) in mice: Inhibition by anpirtoline: A 5-HT1B receptor agonist. Neuropsychopharmacology 27, 171–181 (2002).

17. Amsel, A. & Roussel, J. Motivational properties of frustration: I. Effect on a running response of the addition of frustration to the motivational complex. J. Exp. Psychol. 43, 363–368 (1952).

18. Miller, N. E. & Stevenson, S. S. Agitated behavior of rats during experimental extinction and a curve of spontaneous recovery. undefined 21, 205–231 (1936).

19. Manzo, L., Gómez, M. J., Callejas-Aguilera, J. E., Fernández-Teruel, A., Papini, M. R. & Torres, C. Anti-anxiety self-medication induced by incentive loss in rats. Physiol. Behav. 123, 86–92 (2014).

20. Vindas, M. A., Johansen, I. B., Vela-Avitua, S., Nørstrud, K. S., Aalgaard, M., Braastad, B. O., Höglund, E. & Øverli, Ø. Frustrative reward omission increases aggressive behaviour of inferior fighters. Proc. R. Soc. B Biol. Sci. 281, (2014).

21. Carpenter, R. E., Maruska, K. P., Becker, L. & Fernald, R. D. Social Opportunity Rapidly Regulates Expression of CRF and CRF Receptors in the Brain during Social Ascent of a Teleost Fish, Astatotilapia burtoni. PLoS One 9, e96632 (2014).

22. DeVries, A. C., Glasper, E. R. & Detillion, C. E. Social modulation of stress responses. in Physiology and Behavior vol. 79 399–407 (2003).

23. Zuri, I., Gottreich, A. & Terkel, J. Social stress in neighboring and encountering blind mole-rats (Spalax ehrenbergi). Physiol. Behav. 64, 611–620 (1998).

24. Padgett, D. A., Sheridan, J. F., Dorne, J., Berntson, G. G., Candelora, J. & Glaser, R. Social stress and the reactivation of latent herpes simplex virus type 1. Proc. Natl. Acad. Sci. U. S. A. 95, 7231–7235 (1998).

25. Quan, N., Avitsur, R., Stark, J. L., He, L., Shah, M., Caligiuri, M., Padgett, D. A., Marucha, P. T. & Sheridan, J. F. Social stress increases the susceptibility to endotoxic shock. J. Neuroimmunol. 115, 36–45 (2001).

26. DeVries, A. C., Joh, H. D., Bernard, O., Hattori, K., Hurn, P. D., Traystman, R. J. & Alkayed, N. J. Social stress exacerbates stroke outcome by suppressing Bcl-2 expression. Proc. Natl. Acad. Sci. U. S. A. 98, 11824–11828 (2001).

27. Sugo, N., Hurn, P. D., Morahan, M. B., Hattori, K., Traystman, R. J. & DeVries, A. C. Social stress exacerbates focal cerebral ischemia in mice. Stroke 33, 1660–1664 (2002).

28. Razzoli, M., Nyuyki-Dufe, K., Gurney, A., Erickson, C., McCallum, J., Spielman, N., Marzullo, M., Patricelli, J., Kurata, M., Pope, E. A., Touma, C., Palme, R., Largaespada, D. A., Allison, D. B. & Bartolomucci, A. Social stress shortens lifespan in mice. Aging Cell 17, 1–14 (2018).

29. Pleil, K. E., Rinker, J. A., Lowery-Gionta, E. G., Mazzone, C. M., McCall, N. M., Kendra, A. M., Olson, D. P., Lowell, B. B., Grant, K. A., Thiele, T. E. & Kash, T. L. NPY signaling inhibits extended amygdala CRF neurons to suppress binge alcohol drinking. Nat. Neurosci. 18, 545–552 (2015).

30. Gilpin, N. W., Herman, M. A. & Roberto, M. The Central Amygdala as an Integrative Hub for Anxiety and Alcohol Use Disorders. Biological Psychiatry vol. 77 859–869 (2015).

31. Baik, J. H. Stress and the dopaminergic reward system. Experimental and Molecular Medicine vol. 52 1879–1890 (2020).

32. Ulrich-Lai, Y. M., Christiansen, A. M., Ostrander, M. M., Jones, A. A., Jones, K. R., Choi, D. C., Krause, E. G., Evanson, N. K., Furay, A. R., Davis, J. F., Solomon, M. B., De Kloet, A. D., Tamashiro, K. L., Sakai, R. R., Seeley, R. J., Woods, S. C. & Herman, J. P. Pleasurable behaviors reduce stress via brain reward pathways. Proc. Natl. Acad. Sci. U. S. A. 107, 20529–20534 (2010).

33. Lemos, C., Salti, A., Amaral, I. M., Fontebasso, V., Singewald, N., Dechant, G., Hofer, A. & El Rawas, R. Social interaction reward in rats has anti-stress effects. Addict. Biol. 26, e12878 (2021).

34. Montagud-Romero, S., Blanco-Gandía, M. C., Reguilón, M. D., Ferrer-Pérez, C., Ballestín, R., Miñarro, J. & Rodríguez-Arias, M. Social defeat stress: Mechanisms underlying the increase in rewarding effects of drugs of abuse. European Journal of Neuroscience vol. 48 2948–2970 (2018).

35. Dedic, N., Chen, A. & Deussing, J. M. The CRF Family of Neuropeptides and their Receptors – Mediators of the Central Stress Response. Curr. Mol. Pharmacol. 11, (2017).

36. Deussing, J. M. & Chen, A. The corticotropin-releasing factor family: Physiology of the stress response. Physiological Reviews vol. 98 2225–2286 (2018).

37. Gilpin, N. W. Corticotropin-releasing factor (CRF) and neuropeptide Y (NPY): Effects on inhibitory transmission in central amygdala, and anxiety-& alcohol-related behaviors. Alcohol 46, 329–337 (2012).

38. M. Jiménez & L. Buéno. Inhibitory effects of neuropeptide Y (NPY) on CRF and stress-induced cecal motor response in rats. Life Sci. 47, 205–211 (1990).

39. Sokolowski, M. B. Social interactions in ‘simple’ model systems. Neuron 65, 780–94 (2010).

40. Shohat-Ophir, G., Kaun, K. R., Azanchi, R., Mohammed, H. & Heberlein, U. Sexual deprivation increases ethanol intake in Drosophila. Science 335, 1351–5 (2012).

41. Shankar, S., Chua, J. Y., Tan, K. J., Calvert, M. E. K., Weng, R., Ng, W. C., Mori, K. & Yew, J. Y. The neuropeptide tachykinin is essential for pheromone detection in a gustatory neural circuit. Elife 4, 1–23 (2015).

42. Hoyer, S. C., Eckart, A., Herrel, A., Zars, T., Fischer, S. A., Hardie, S. L. & Heisenberg, M. Octopamine in Male Aggression of Drosophila. Curr. Biol. 18, 159–167 (2008).

43. Liu, W., Liang, X., Gong, J., Yang, Z., Zhang, Y. H., Zhang, J. X. & Rao, Y. Social regulation of aggression by pheromonal activation of Or65a olfactory neurons in Drosophila. Nat. Neurosci. 14, 896–902 (2011).

44. Wang, L., Dankert, H., Perona, P. & Anderson, D. J. A common genetic target for environmental and heritable influences on aggressiveness in Drosophila. Proc. Natl. Acad. Sci. U. S. A. 105, 5657–63 (2008).

45. Ejima, A., Smith, B. P. C., Lucas, C., van der Goes van Naters, W., Miller, C. J., Carlson, J. R., Levine, J. D. & Griffith, L. C. Generalization of courtship learning in Drosophila is mediated by cis-vaccenyl acetate. Curr. Biol. 17, 599–605 (2007).

46. Certel, S. J., Savella, M. G., Schlegel, D. C. F. F. & Kravitz, E. A. Modulation of Drosophila male behavioral choice. PNAS 104, 4706–4711 (2007).

47. Omesi, L., Levi, M., Bentzur, A., Kim, Y.-K., Ben-Shaanan, S., Azanchi, R. & Shohat-Ophir, G. Sexual deprivation modulates social interaction and reproductive physiology. bioRxiv 2021.04.27.441612 (2021).

48. Kim, W. J., Jan, L. Y. & Jan, Y. N. A PDF/NPF neuropeptide signaling circuitry of male Drosophila melanogaster controls rival-induced prolonged mating. Neuron 80, 1190–1205 (2013).

49. Kim, W. J., Jan, L. Y. & Jan, Y. N. Contribution of visual and circadian neural circuits to memory for prolonged mating induced by rivals. Nat. Neurosci. 15, 876–83 (2012).

50. Bretman, A., Fricke, C., Hetherington, P., Stone, R. & Chapman, T. Exposure to rivals and plastic responses to sperm competition in Drosophila melanogaster. Behav. Ecol. 21, 317–321 (2010).

51. Asahina, K., Watanabe, K., Duistermars, B. J., Hoopfer, E., González, C. R., Eyjólfsdóttir, E. A., Perona, P. & Anderson, D. J. Tachykinin-expressing neurons control male-specific aggressive arousal in drosophila. Cell 156, 221–235 (2014).

52. Bentzur, A., Shmueli, A., Omesi, L., Ryvkin, J., Knapp, J. M., Parnas, M., Davis, F. P. & Shohat-Ophir, G. Odorant binding protein 69a connects social interaction to modulation of social responsiveness in Drosophila. PLoS Genet. 14, 1–23 (2018).

53. Catalano, J., Mei, N., Azanchi, R., Song, S., Blackwater, T., Heberlein, U. & Kaun, K. Behavioral features of motivated response to alcohol in Drosophila. bioRxiv 2020.02.17.953026 (2020).

54. Ryvkin, J., Bentzur, A., Zer-krispil, S. & Shohat-Ophir, G. Mechanisms Underlying the Risk to Develop Drug Addiction, Insights From Studies in Drosophila melanogaster. Front. Physiol. 9, 327 (2018).

55. Dvořáček, J. & Kodrík, D. Drosophila reward system – A summary of current knowledge. Neuroscience and Biobehavioral Reviews vol. 123 301–319 (2021).

56. Xu, J., Li, M. & Shen, P. A G-protein-coupled neuropeptide Y-like receptor suppresses behavioral and sensory response to multiple stressful stimuli in Drosophila. J. Neurosci. 30, 2504–12 (2010).

57. Walker, R. J., Papaioannou, S. & Holden-Dye, L. A review of FMRFamide-and RFamide-like peptides in metazoa. Invertebrate Neuroscience vol. 9 111–153 (2009).

58. Cardoso, J. C. R., Félix, R. C., Bergqvist, C. A. & Larhammar, D. New insights into the evolution of vertebrate CRH (corticotropin-releasing hormone) and invertebrate DH44 (diuretic hormone 44) receptors in metazoans. Gen. Comp. Endocrinol. 209, 162–170 (2014).

59. Zhang, S. X., Rogulja, D. & Crickmore, M. A. Recurrent Circuitry Sustains Drosophila Courtship Drive While Priming Itself for Satiety. Curr. Biol. 29, 3216–3228.e9 (2019).

60. Ryvkin, J., Bentzur, A., Shmueli, A., Tannenbaum, M., Shallom, O., Dokarker, S., C., B. J. I., Levi, M. & Shohat-Ophir, G. Transcriptome analysis of NPFR neurons reveals a connection between proteome diversity and social behavior. Front. Behav. Neurosci. 15, 35 (2021).

61. Krashes, M. J., DasGupta, S., Vreede, A., White, B., Douglas Armstrong, J., Waddell, S., Armstrong, J. D. & Waddell, S. A neural circuit mechanism integrating motivational state with memory expression in Drosophila. Cell 139, 416–27 (2009).

62. Dierick, H. A. & Greenspan, R. J. Serotonin and neuropeptide F have opposite modulatory effects on fly aggression. Nat. Genet. 39, 678–682 (2007).

63. Lingo, P. R., Zhao, Z. & Shen, P. Co-regulation of cold-resistant food acquisition by insulin-and neuropeptide Y-like systems in Drosophila melanogaster. Neuroscience 148, 371–374 (2007).

64. Beshel, J. & Zhong, Y. Graded Encoding of Food Odor Value in the Drosophila Brain. J. Neurosci. 33, 15693–15704 (2013).

65. Kacsoh, B. Z., Lynch, Z. R., Mortimer, N. T. & Schlenke, T. A. Fruit flies medicate offspring after seeing parasites. Science 339, 947–50 (2013).

66. Johnson, E. C., Garczynski, S. F., Park, D., Crim, J. W., Nässel, D. R. & Taghert, P. H. Identification and characterization of a G protein-coupled receptor for the neuropeptide proctolin in Drosophila melanogaster. Proc. Natl. Acad. Sci. U. S. A. 100, 6198–6203 (2003).

67. Lee, G., Bahn, J. H. & Park, J. H. Sex-and clock-controlled expression of the neuropeptide F gene in Drosophila. Proc. Natl. Acad. Sci. U. S. A. 103, 12580–12585 (2006).

68. Wen, T., Parrish, C. a, Xu, D., Wu, Q. & Shen, P. Drosophila neuropeptide F and its receptor, NPFR1, define a signaling pathway that acutely modulates alcohol sensitivity. Proc. Natl. Acad. Sci. U. S. A. 102, 2141–6 (2005).

69. Kim, Y. K., Saver, M., Simon, J., Kent, C. F., Shao, L., Eddison, M., Agrawal, P., Texada, M., Truman, J. W. & Heberlein, U. Repetitive aggressive encounters generate a long-lasting internal state in Drosophila melanogaster males. Proc. Natl. Acad. Sci. U. S. A. 115, 1099–1104 (2018).

70. Kalra, S. P. & Kalra, P. S. To subjugate NPY is to improve the quality of life and live longer. Peptides 28, 413–418 (2007).

71. Kalra, S. Global Life-Long Health Benefits of Repression of Hypothalamic NPY System by Central Leptin Gene Therapy. Curr. Top. Med. Chem. 7, 1675–1681 (2007).

72. Chiba, T., Tamashiro, Y., Park, D., Kusudo, T., Fujie, R., Komatsu, T., Kim, S. E., Park, S., Hayashi, H., Mori, R., Yamashita, H., Chung, H. Y. & Shimokawa, I. A key role for neuropeptide y in lifespan extension and cancer suppression via dietary restriction. Sci. Rep. 4, 1–10 (2014).

73. Michalkiewicz, M., Knestaut, K. M., Bytchkova, E. Y. & Michalkiewicz, T. Hypotension and reduced catecholamines in neuropeptide Y transgenic rats. Hypertension 41, 1056–1062 (2003).

74. SJ, A. & H, H. NPY family of peptides in neurobiology, cardiovascular and metabolic disorders: from genes to therapeutics. in Springer Science & Business Media (eds. Z, Z. & GZ, F.) vol. 95 113–223 (Birkhäuser Basel, 2006).

75. Gendron, C. M., Kuo, T. H., Harvanek, Z. M., Chung, B. Y., Yew, J. Y., Dierick, H. A. & Pletcher, S. D. Drosophila life span and physiology are modulated by sexual perception and reward. Science (80-.). 343, 544–548 (2014).

76. Chung, B. Y., Ro, J., Hutter, S. A., Miller, K. M., Guduguntla, L. S., Kondo, S. & Pletcher, S. D. Drosophila Neuropeptide F Signaling Independently Regulates Feeding and Sleep-Wake Behavior. Cell Rep. 19, 2441–2450 (2017).

77. Zer-Krispil, S., Zak, H., Shao, L., Ben-Shaanan, S., Tordjman, L., Bentzur, A., Shmueli, A. & Shohat-Ophir, G. Ejaculation Induced by the Activation of Crz Neurons Is Rewarding to Drosophila Males. Curr. Biol. 28, 1445–1452.e3 (2018).

78. Gao, C., Guo, C., Peng, Q., Cao, J., Shohat-Ophir, G., Liu, D. & Pan, Y. Sex and Death: Identification of Feedback Neuromodulation Balancing Reproduction and Survival. Neurosci. Bull. 36, 1429–1440 (2020).

79. Ejima, A., Smith, B. P. C., Lucas, C., Levine, J. D. & Griffith, L. C. Sequential learning of pheromonal cues modulates memory consolidation in trainer-specific associative courtship conditioning. Curr. Biol. 15, 194–206 (2005).

80. Siegel, R. W. & Hall, J. C. Conditioned responses in courtship behavior of normal and mutant *Drosophila*. Proc. Natl. Acad. Sci. U. S. A. 76, 3430–3434 (1979).

81. Mehren, J. E., Ejima, A. & Griffith, L. C. Unconventional sex: fresh approaches to courtship learning. Curr. Opin. Neurobiol. 14, 745–50 (2004).

82. Morley, J. E. & Levine, A. S. Corticotrophin releasing factor, grooming and ingestive behavior. Life Sci. 31, 1459–1464 (1982).

83. Moyaho, A. & Valencia, J. Grooming and yawning trace adjustment to unfamiliar environments in laboratory Sprague-Dawley rats (Rattus norvegicus). J. Comp. Psychol. 116, 263–269 (2002).

84. Troisi, A. Displacement activities as a behavioral measure of stress in nonhuman primates and human subjects. Stress 5, 47–54 (2002).

85. Gould, T. D., Dao, D. T. & Kovacsics, C. E. Mood and Anxiety Related Phenotypes in Mice. Neuromethods 42, 1–20 (2009).

86. Harvanek, Z. M., Lyu, Y., Gendron, C. M., Johnson, J. C., Kondo, S., Promislow, D. E. L. & Pletcher, S. D. Perceptive costs of reproduction drive ageing and physiology in male Drosophila. Nat. Ecol. Evol. 1, 1–15 (2017).

87. Henry, G. L., Davis, F. P., Picard, S. & Eddy, S. R. Cell type-specific genomics of Drosophila neurons. Nucleic Acids Res. 40, 9691–704 (2012).

88. Okamoto, N. & Nishimura, T. Signaling from Glia and Cholinergic Neurons Controls Nutrient-Dependent Production of an Insulin-like Peptide for Drosophila Body Growth. Dev. Cell 35, 295–310 (2015).

89. Orgad, S., Rosenfeld, G., Greenspan, R. J. & Segal, D. courtless, The Drosophila UBC7 homolog, is involved in male courtship behavior and spermatogenesis. Genetics 155, 1267–1280 (2000).

90. Post, S., Karashchuk, G., Wade, J. D., Sajid, W., De Meyts, P. & Tatar, M. Drosophila insulin-like peptides DILP2 and DILP5 differentially stimulate cell signaling and glycogen phosphorylase to regulate longevity. Front. Endocrinol. (Lausanne). 9, (2018).

91. Nässel, D. R. Substrates for neuronal cotransmission with neuropeptides and small molecule neurotransmitters in drosophila. Frontiers in Cellular Neuroscience vol. 12 (2018).

92. Kannan, K. & Fridell, Y.-W. C. Functional implications of Drosophila insulin-like peptides in metabolism, aging, and dietary restriction. Front. Physiol. 4, (2013).

93. Manière, G., Ziegler, A. B., Geillon, F., Featherstone, D. E. & Grosjean, Y. Direct Sensing of Nutrients via a LAT1-like Transporter in Drosophila Insulin-Producing Cells. Cell Rep. 17, 137–148 (2016).

94. Scharf, M. E., Scharf, D. W., Bennett, G. W. & Pittendrigh, B. R. Catalytic activity and expression of two flavin-containing monooxygenases from Drosophila melanogaster. Arch. Insect Biochem. Physiol. 57, 28–39 (2004).

95. Andres, A. J., Fletcher, J. C., Karim, F. D. & Thummel, C. S. Molecular Analysis of the Initiation of Insect Metamorphosis: A Comparative Study of Drosophila Ecdysteroid-Regulated Transcription. Dev. Biol. 160, 388–404 (1993).

96. Maciejczyk, M., Żebrowska, E. & Chabowski, A. Insulin resistance and oxidative stress in the brain: What’s new? Int. J. Mol. Sci. 20, 874 (2019).

97. Zhang, X., Beaulieu, J. M., Sotnikova, T. D., Gainetdinov, R. R. & Caron, M. G. Tryptophan hydroxylase-2 controls brain synthesis. Science (80-.). 305, 217 (2004).

98. Mahony, S. M. O., Clarke, G., Borre, Y. E., Dinan, T. G. & Cryan, J. F. Serotonin, tryptophan metabolism and the brain-gut-microbiome axis. Behav. Brain Res. 277, 32–48 (2015).

99. Platten, M. & Fallarino, F. Tryptophan metabolism as a common therapeutic target in cancer, neurodegeneration and beyond. Nat. Rev. Drug Discov. 18, (2019).

100. P.D, L. Tryptophan availability and serotonin synthesis. in Paoletti R., Vanhoutte P.M., Brunello N., Maggi F.M. (eds) Serotonin vol. 46 143–156 (Springer, Dordrecht, 1987).

101. Broughton, S., Alic, N., Slack, C., Bass, T., Ikeya, T., Vinti, G., Tommasi, A. M., Driege, Y., Hafen, E. & Partridge, L. Reduction of DILP2 in Drosophila triages a metabolic phenotype from lifespan revealing redundancy and compensation among DILPs. PLoS One 3, 3–11 (2008).

102. Belgacem, Y. H. & Martin, J. R. Disruption of insulin pathways alters trehalose level and abolishes sexual dimorphism in locomotor activity in Drosophila. J. Neurobiol. 66, 19–32 (2006).

103. Rulifson, E. J., Kim, S. K. & Nusse, R. Ablation of insulin-producing neurons in files: Growth and diabetic phenotypes. Science (80-.). 296, 1118–1120 (2002).

104. Reyes-DelaTorre, A., Pena-Rangel, M. T. & Riesgo-Escovar, J. R. Carbohydrates – Comprehensive Studies on Glycobiology and Glycotechnology. in inTech (ed. Chang, C.-F.) 317–338 (BoD – Books on Demand, 2012).

105. Hwangbo, D. S., Garsham, B., Tu, M. P., Palmer, M. & Tatar, M. Drosophila dFOXO controls lifespan and regulates insulin signalling in brain and fat body. Nature 429, 562–566 (2004).

106. Broughton, S. J., Piper, M. D. W., Ikeya, T., Bass, T. M., Jacobson, J., Driege, Y., Martinez, P., Hafen, E., Withers, D. J., Leevers, S. J. & Partridge, L. Longer lifespan, altered metabolism, and stress resistance in Drosophila from ablation of cells making insulin-like ligands. Proc. Natl. Acad. Sci. U. S. A. 102, 3105–3110 (2005).

107. Piper, M. D. W., Selman, C., McElwee, J. J. & Partridge, L. Separating cause from effect: How does insulin/IGF signalling control lifespan in worms, flies and mice? J. Intern. Med. 263, 179–191 (2008).

108. Giannakou, M. E. & Partridge, L. Role of insulin-like signalling in Drosophila lifespan. Trends Biochem. Sci. 32, 180–188 (2007).

109. De Rosa, M. J., Veuthey, T., Florman, J., Grant, J., Blanco, M. G., Andersen, N., Donnelly, J., Rayes, D. & Alkema, M. J. The flight response impairs cytoprotective mechanisms by activating the insulin pathway. Nature 573, 135–138 (2019).

110. Kim, S. Y. & Webb, A. E. Neuronal functions of FOXO/DAF-16. Nutr. Heal. Aging 4, 113–126 (2017).

111. Lehtinen, M. K., Yuan, Z., Boag, P. R., Yang, Y., Villén, J., Becker, E. B. E., DiBacco, S., de la Iglesia, N., Gygi, S., Blackwell, T. K. & Bonni, A. A Conserved MST-FOXO Signaling Pathway Mediates Oxidative-Stress Responses and Extends Life Span. Cell 125, 987–1001 (2006).

112. Bai, H., Kang, P. & Tatar, M. Drosophila insulin-like peptide-6 (dilp6) expression from fat body extends lifespan and represses secretion of Drosophila insulin-like peptide-2 from the brain. Aging Cell 11, 978–985 (2012).

113. Wang, B., Moya, N., Niessen, S., Hoover, H., Mihaylova, M. M., Shaw, R. J., Yates, J. R., Fischer, W. H., Thomas, J. B. & Montminy, M. A hormone-dependent module regulating energy balance. Cell 145, 596–606 (2011).

114. Vihervaara, T. & Puig, O. dFOXO Regulates Transcription of a Drosophila Acid Lipase. J. Mol. Biol. 376, 1215–1223 (2008).

115. Cao, J., Ni, J., Ma, W., Shi, V., Milla, L. A., Park, S., Spletter, M. L., Tang, S., Zhang, J., Wei, X., Kim, S. K. & Scott, M. P. Insight into insulin secretion from transcriptome and genetic analysis of insulin-producing cells of Drosophila. Genetics 197, 175–192 (2014).

116. Bitran, M., Torres, G., Fournier, A., Pierre, S. S. & Pablo Huidobro-Toro, J. Age and castration modulate the inhibitory action of neuropeptide Y on neurotransmission In the rat vas deferens. Eur. J. Pharmacol. 203, 267–274 (1991).

117. Rajpara, S. M., Garcia, P. D., Roberts, R., Eliassen, J. C., Owens, D. F., Maltby, D., Myers, R. M. & Mayeri, E. Identification and molecular cloning of a neuropeptide y homolog that produces prolonged inhibition in aplysia neurons. Neuron 9, 505–513 (1992).

118. Browning, K. N. & Travagli, R. A. Neuropeptide Y and Peptide YY Inhibit Excitatory Synaptic Transmission in the Rat Dorsal Motor Nucleus of the Vagus. J. Physiol. 549, 775–785 (2003).

119. Bentzur, A., Ben-Shaanan, S., Benishou, J., Costi, E., Levi, M., Ilany, A. & Shohat-Ophir, G. Early Life Experience Shapes Male Behavior and Social Networks in Drosophila. Curr. Biol. 31, 486–501.e3 (2020).

120. Shao, L., Saver, M., Chung, P., Ren, Q., Lee, T., Kent, C. F. & Heberlein, U. Dissection of the Drosophila neuropeptide F circuit using a high-throughput two-choice assay. Proc. Natl. Acad. Sci. U. S. A. 114, E8091–E8099 (2017).

121. Pimentel, E., Vidal, L. M., Cruces, M. P. & Janczur, M. K. Action of protoporphyrin-IX (PP-IX) in the lifespan of Drosophila melanogaster deficient in endogenous antioxidants, Sod and Cat. Open J. Anim. Sci. 03, 1–7 (2013).

122. Williams, M., Krootjes, B. B. H., van Steveninck, J. & van Der Zee, J. The pro-and antioxidant properties of protoporphyrin IX. Biochim. Biophys. Acta (BBA)/Lipids Lipid Metab. 1211, 310–316 (1994).

123. Afonso, S., Vanore, G. & Batlle, A. Protoporphyrin IX and oxidative stress. Free Radic. Res. 31, 161–170 (1999).

124. Sachar, M., Anderson, K. E. & Ma, X. Protoporphyrin IX: The good, the bad, and the ugly. J. Pharmacol. Exp. Ther. 356, 267–275 (2016).

125. Kikuchi, G., Yoshida, T. & Noguchi, M. Heme oxygenase and heme degradation. Biochemical and Biophysical Research Communications vol. 338 558–567 (2005).

126. Abaquita, T. A. L., Damulewicz, M., Bhattacharya, D. & Pyza, E. Regulation of heme oxygenase and its cross-talks with apoptosis and autophagy under different conditions in drosophila. Antioxidants 10, 1716 (2021).

127. Nässel, D. R., Zandawala, M., Kawada, T. & Satake, H. Tachykinins: Neuropeptides That Are Ancient, Diverse, Widespread and Functionally Pleiotropic. Front. Neurosci. 13, 1262 (2019).

128. Winther, Å. M. E., Siviter, R. J., Isaac, R. E., Predel, R. & Nässel, D. R. Neuronal expression of tachykinin-related peptides and gene transcript during postembryonic development of Drosophila. J. Comp. Neurol. 464, 180–196 (2003).

129. Davis, F. P., Nern, A., Picard, S., Reiser, M. B., Rubin, G. M., Eddy, S. R. & Henry, G. L. A genetic, genomic, and computational resource for exploring neural circuit function. Elife 9, 1–40 (2020).

130. Alon, S., Goodwin, D. R., Sinha, A., Wassie, A. T., Chen, F., Daugharthy, E. R., Bando, Y., Kajita, A., Xue, A. G., Marrett, K., Prior, R., Cui, Y., Payne, A. C., Yao, C.-C., Suk, H.-J., Wang, R., Yu, C.-C. (Jay), … Boyden, E. S. Expansion sequencing: Spatially precise in situ transcriptomics in intact biological systems. Science (80-.). 371, eaax2656 (2021).

131. Keeney, A., Jessop, D. S., Harbuz, M. S., Marsden, C. A., Hogg, S. & Blackburn-Munro, R. E. Differential Effects of Acute and Chronic Social Defeat Stress on Hypothalamic-Pituitary-Adrenal Axis Function and Hippocampal Serotonin Release in Mice. J. Neuroendocrinol. 18, 330–338 (2006).

132. Newman, E. L., Leonard, M. Z., Arena, D. T., de Almeida, R. M. M. & Miczek, K. A. Social defeat stress and escalation of cocaine and alcohol consumption: Focus on CRF. Neurobiology of Stress vol. 9 151–165 (2018).

133. Boyson, C. O., Holly, E. N., Shimamoto, A., Albrechet-Souza, L., Weiner, L. A., DeBold, J. F. & Miczek, K. A. Social Stress and CRF-Dopamine Interactions in the VTA: Role in Long-Term Escalation of Cocaine Self-Administration. J. Neurosci. 34, 6659–6667 (2014).

134. Boyson, C. O., Miguel, T. T., Quadros, I. M., DeBold, J. F. & Miczek, K. A. Prevention of social stress-escalated cocaine self-administration by CRF-R1 antagonist in the rat VTA. Psychopharmacology (Berl). 218, 257–269 (2011).

135. Hwa, L. S., Holly, E. N., Debold, J. F. & Miczek, K. A. Social stress-escalated intermittent alcohol drinking: modulation by CRF-R1 in the ventral tegmental area and accumbal dopamine in mice. doi:10.1007/s00213-015-4144-2.

136. Tovar-Díaz, J., Pomrenze, M. B., Kan, R., Pahlavan, B. & Morikawa, H. Cooperative CRF and α1 Adrenergic Signaling in the VTA Promotes NMDA Plasticity and Drives Social Stress Enhancement of Cocaine Conditioning. Cell Rep. 22, 2756–2766 (2018).

137. Martin, M. Cutadapt removes adapter sequences from high-throughput sequencing reads. EMBnet.journal 17, 10 (2011).

138. Dobin, A., Davis, C. A., Schlesinger, F., Drenkow, J., Zaleski, C., Jha, S., Batut, P., Chaisson, M. & Gingeras, T. R. STAR: Ultrafast universal RNA-seq aligner. Bioinformatics 29, 15–21 (2013).

139. Anders, S., Pyl, P. T. & Huber, W. HTSeq-A Python framework to work with high-throughput sequencing data. Bioinformatics 31, 166–169 (2015).

140. Liao, Y., Smyth, G. & Shi, W. featureCounts: an efficient general purpose program for assigning sequence reads to genomic features. Bioinformatics 30, 923–930 (2014).

141. Love, M. I., Huber, W. & Anders, S. Moderated estimation of fold change and dispersion for RNA-seq data with DESeq2. Genome Biol. 15, 550 (2014).

142. Branson, K., Robie, A. A., Bender, J., Perona, P. & Dickinson, M. H. High-throughput ethomics in large groups of Drosophila. Nat. Methods 6, 451–457 (2009).

143. Kabra, M., Robie, A. A., Rivera-Alba, M., Branson, S. & Branson, K. JAABA: Interactive machine learning for automatic annotation of animal behavior. Nat. Methods 10, 64–67 (2013).

144. Tennessen, J. M., Barry, W. E., Cox, J. & Thummel, C. S. Methods for studying metabolism in Drosophila. Methods 68, 105–115 (2014).

145. MacKay, G. M., Zheng, L., Van Den Broek, N. J. F. & Gottlieb, E. Analysis of Cell Metabolism Using LC-MS and Isotope Tracers. Methods Enzymol. 561, 171–196 (2015).

146. Pietzke, M. & Vazquez, A. Metabolite AutoPlotter – an application to process and visualise metabolite data in the web browser. Cancer Metab. 2020 81 8, 1–11 (2020).

147. Thomas, P. D., Kejariwal, A., Guo, N., Mi, H., Campbell, M. J., Muruganujan, A. & Lazareva-Ulitsky, B. Applications for protein sequence-function evolution data: mRNA/protein expression analysis and coding SNP scoring tools. Nucleic Acids Res. 34, W645–W650 (2006).

148. Mi, H., Muruganujan, A., Ebert, D., Huang, X. & Thomas, P. D. PANTHER version 14: More genomes, a new PANTHER GO-slim and improvements in enrichment analysis tools. Nucleic Acids Res. 47, D419–D426 (2019).

149. Bouliotis, G. & Billingham, L. Crossing survival curves: alternatives to the log-rank test. Trials 12, 6215 (2011).

